# Cell identity and activation state determine the innate immune response to nanoplastics

**DOI:** 10.64898/2026.11.16.767123

**Authors:** Aitana Ignes-Romeu, Yoanna I. Ivanova, Justin M. Owens, Eric L. Ginter, Jacob W. Burleson, Margaret Radke, Hannah K. Weppner, Lea P. Hibbard, Laurel E. Hind

## Abstract

Autoimmune disease is on the rise, and while no direct connection has been made, evidence suggests environmental toxins such as nanoplastics may be contributing to the increased incidence. A key factor in the onset of autoimmunity is inflammation; yet how nanoplastics impact innate immune function is unclear. Here, we exposed primary human neutrophils, macrophages, and endothelial cells to biocorona-coated polystyrene nanoplastics at physiologically relevant concentrations. All cells internalized nanoplastics without viability loss, but functional consequences diverged. Neutrophils increased myeloperoxidase release as nanoplastic concentration increased. Endothelial cells reduced leukocyte-recruiting mediator secretion and had decreased neutrophil adhesion following inflammatory stimulation. M1 macrophages broadly upregulated inflammatory cytokine secretion (including TNF-α and IL-6), whereas M0 and M2 macrophages exhibited no significant secretory changes. Collectively, these findings indicate physiologically relevant nanoplastic concentrations do not universally increase inflammation but rather dysregulate innate immune populations in a manner that could contribute to autoimmunity.

## Introduction

Autoimmune disorders are on the rise, now affecting roughly one in ten people. The cause of this increase has not been established; however, socioeconomic, seasonal, and regional gradients in incidence suggest that environmental factors are contributors^1^. The establishment of autoimmunity requires a genetic predisposition, an environmental exposure, and immune dysregulation^2^. Environmental exposure to toxins can lead to chronic, low-grade inflammation caused by sustained mediator secretion, tissue damage, and inefficient clearance of debris that dysregulates the immune response^3^. Nanoplastics are a candidate environmental toxin as they have reached measurable levels in human blood and tissue^4–7^, are internalized by immune cells but cannot be degraded^8^, and are consistently reported to be inflammatory across cell types and models^9–11^. Attributing a specific environmental toxin to autoimmune disease requires a long chain of evidence, and the first link in that chain is the effect exposure has on the behavior of innate immune cells. This study attempts to answer that question by investigating how nanoplastic exposure at concentrations found in tissues and blood alters innate immune associated cell function.

Across cell types and experimental approaches, the most consistently documented response to NPs is inflammatory, including increased oxidative stress, IL-1β and IL-18 release, genotoxicity, and reduced cell viability at higher concentrations^12^. However, many of the NP concentrations used are supraphysiological so their effects could be overstated^4,12^. Additionally, surface properties vary greatly; surface properties govern how a particle is recognized and internalize^13,14^, and upon entering the body the surface of NPs become coated in a biocorona with different surface properties than bare polymer^15^. Despite considerable variability between studies, there is consensus that NPs affect the immune response and human health. That variability nonetheless makes the literature difficult to reconcile, because studies differ in cell type, NP polymer, concentration, exposure time, and surface chemistry, and each of these variables can affect the cellular response. In our study, we sought to use multiple primary human cell types exposed to identical NPs under physiologically relevant concentrations to increase the applicability of our findings.

Understanding the effects of NPs on the innate immune response requires study in the context of the bloodstream, which is both a route of NP distribution and the compartment through which innate immune cells travel^4,11^. During an inflammatory response, these cells act in sequence. In the vasculature, endothelial cells (ECs) sense damage or infection, and the junctions maintaining the barrier between blood and tissue are disrupted^16^. ECs then mediate leukocyte migration through adhesion molecules such as ICAM-1 and through secreted chemokines^17,18^. Neutrophils arrive first among leukocytes and release preformed granule contents, of which myeloperoxidase (MPO) is the most abundant^19,20^. Monocytes follow and differentiate into macrophages, which, together with tissue-resident macrophages, regulate the local response through their activation state: resting (M0), classically activated (M1) and alternatively activated (M2) macrophages differ in both engulfment of particles and mediator release^21,22^.

There is experimental evidence that NPs influence the function of cells critical to the innate immune response, yet a comprehensive understanding of how each cell is affected under the same NP conditions has not been developed. Understanding the alteration in innate immune function under relevant NP conditions is the first step in understanding their role in the development of autoimmunity. Here, we compare the response of primary human neutrophils, endothelial cells, and macrophages following exposure to identically treated NPs. The particles carried a biocorona formed in human plasma and were applied at concentrations reported in human blood^4^. Across the three cell types, we measured NP internalization, viability, and inflammatory function. We find that NPs were internalized by all three cell types without reducing viability, but that their functional consequences depended on cell type and activation state. NP exposure reduced intracellular MPO in neutrophils at the highest concentration, reduced endothelial mediator secretion and neutrophil adhesion to an inflamed endothelium, and increased cytokine secretion by classically activated macrophages without altering their phagocytosis.

## Results

### Incubation in human plasma forms a biocorona on NPs

NPs enter the body primarily through ingestion and inhalation, and particles that cross the intestinal or airway epithelium can reach the bloodstream^23^. Within minutes of entering the bloodstream, NPs are coated with plasma proteins, lipids, and other biomolecules, forming a biocorona^13,14^. The biocorona, rather than the bare polymer, interacts with cells and influences how NPs are recognized and internalized^13,14^. To replicate this process experimentally, commercial polystyrene NPs were incubated in human plasma for 10 min or 24 h (Fig. 1A). Biocorona formation was visualized by TEM (Fig. 1B), and particle size and surface charge were measured by DLS and zeta-potential analysis (Fig. 1C and D).

**Figure 1.**
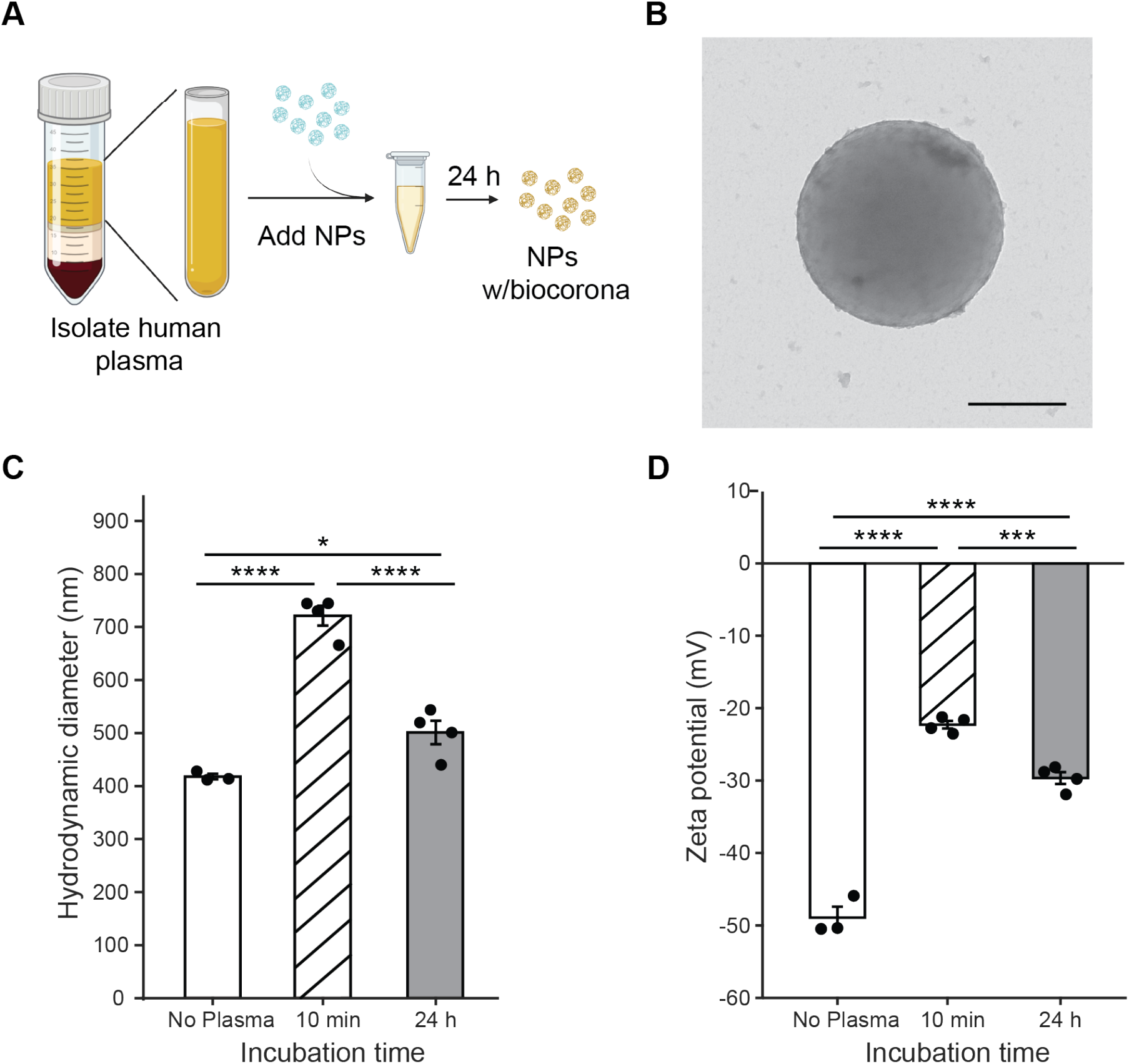
Nanoplastic incubation in human plasma creates a biocorona. **A** Schematic of human plasma isolation and biocorona formation on the surface of polystyrene NPs. **B** TEM image showing biocorona coating on the surface of unweathered NPs. Scale bar, 100 nm. **C** NP hydrodynamic diameter. **D** NP zeta potential. (**C** and **D**) One-way ANOVA with Tukey’s post-hoc comparisons. Plasma incubated samples had N = 4, n=1 replicates; each biological replicate used plasma from a different donor. No plasma control has N=3, n=1 replicates. Bars show the mean ± s.e.m. with individual biological replicates overlaid.*P < 0.05, ***P < 0.001, ****P < 0.0001. N, biological replicates (donors); n, technical replicates per biological replicate and condition.

After 10 min, the hydrodynamic diameter of plasma-coated NPs was larger than that of the no-plasma control (721.3 versus 417.9 nm). Plasma-coated NPs were smaller at 24h than at 10 min but remained larger than NPs without plasma (501.1 nm; Fig. 1C). The change in size between the 10-minute incubation and 24h incubation indicates a transition from a loosely bound soft corona, which forms first, to a tightly bound hard corona. NPs incubated in plasma also had a zeta potential closer to zero than control NPs (−22.3 and −29.7 versus −48.9 mV after 10 min and 24 h; Fig. 1D), consistent with adsorbed proteins masking the negatively charged particle surface.

Fluorescent NPs used to quantify uptake showed the same evidence of a biocorona. After 24 h in plasma, NP hydrodynamic diameter was larger than without plasma (582.1 versus 505.5 nm) and their zeta potential has a small decrease in magnitude (−27.6 versus −23.1 mV), reaching values similar to those of non-fluorescent NPs incubated for 24 h (Fig. S1). Together, these results show incubation in plasma changed the size and surface charge of both NP types within 24 h. All NPs used in subsequent experiments were incubated in human plasma for 24 h before use.

### NPs are internalized by endothelial cells, neutrophils and macrophages without reducing viability

Cells cannot degrade polystyrene, so internalized NPs are retained and accumulate within the cell, where increasing levels of NP could alter cellular function^8^. Primary human endothelial cells, neutrophils and macrophages were exposed to 0, 1, 10 or 50 µg/mL plasma-coated NPs. Exposure times matched the lifespan and function of each cell type: 1 or 7 days of daily exposure for human brain microvascular (hBMEC) and umbilical vein (HUVEC) endothelial cells, to model sustained contact with circulating NPs; 2 h for neutrophils, which only survive for hours after isolation; and 24 h for monocyte-derived macrophages polarized to the M0, M1 or M2 state.

One concern with NP accumulation is reduced cell viability, which could confound functional measurements; therefore, we first measured viability in each cell type. Endothelial viability was unaffected by NP exposure over 7 days in either hBMEC or HUVEC (Fig. 2A). Neutrophil viability was also unaffected by NP exposure. While viability decreased over time, this is expected for isolated primary human neutrophils and did not change with NP exposure. In contrast, the positive control, PMA, which induces neutrophil death reduced viability to zero by 4 h (Fig. 2B and Fig. S2). Macrophage viability was also unaffected by NP exposure in all three phenotypes (Fig. 2C). Together, these results show NP exposure at all tested concentrations does not reduce viability in any cell type over the experimental window.

**Figure 2.**
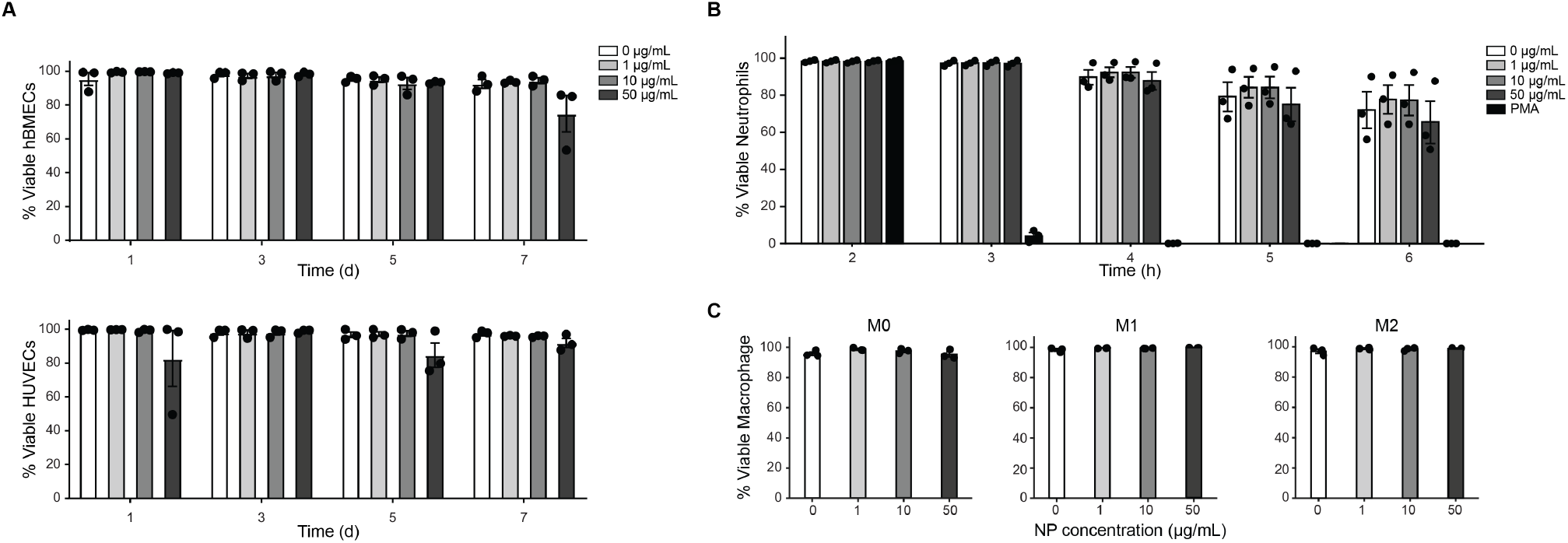
NP exposure does not reduce viability of endothelial cells, neutrophils, or macrophages. **A** hBMEC and HUVEC viability over 7 days of daily NP exposure. Two-way ANOVA with Tukey’s post-hoc comparisons; for hBMECs .N = 3, n = 9. **B** Neutrophil viability over 6 h following blood draw, with PMA as a positive control for cell death. N = 3, n = 6. **C** M0, M1 and M2 macrophage viability after 24 h of NP exposure. N = 3, n = 3. (**B** and **C**) One-way ANOVA with Tukey’s post-hoc comparisons. Comparisons between PMA and NP conditions are not shown for clarity. Bars show the mean ± s.e.m. with individual biological replicates overlaid. N, biological replicates (donors); n, technical replicates (fields of view) per biological replicate and condition.

Endothelial cells function as a continuous monolayer; therefore, we also measured cellular area coverage and the number of nuclei, which report cell loss, spreading, and proliferation. Neither value was affected by NP exposure after 1 day (Fig. S3A and B). After 7 days, area coverage was higher for cells exposed to 50 µg/mL in both EC types and nuclei number was higher in HUVEC exposed to 10 µg/mL (Fig. S3A and B), with greater variation between biological replicates than after 1 day. Because a monolayer can remain intact while its cells become less active, we also measured metabolic activity by alamarBlue. Metabolic activity was unaffected by NP exposure after 1 day in either cell type. At 7 days, HUVECs continued to show no difference in metabolic activity. However, hBMECs showed decreased metabolic activity when exposed to 50 µg/mL compared to anvuntreated control (Fig. S3D). Given our findings that after 7 days of NP exposure, area coverage for cells exposed to 50 µg/mL was higher in both EC types and that metabolic activity at 50 µg/mL was lower in hBMEC, functional assays at this timepoint were limited to concentrations up to 10 µg/mL, to compare secretion and adhesion across equivalent cellular monolayers.

To determine whether NPs were entering the cells or remaining bound to their surface, we imaged each cell type in three dimensions. Fluorescence measured in a single optical plane cannot distinguish internalized particles from particles bound to the cell surface. We therefore reconstructed three-dimensional images of cells following NP incubation and, for phagocytes, phagocytosis of zymosan particles (Fig. S4). We classified each NP surface, and in phagocytes each zymosan surface, as intracellular or extracellular by its position relative to the plasma membrane. NPs were intracellular in ECs (Fig. 3A), neutrophils (Fig. 3B), and macrophages (Fig. 3C)

**Figure 3.**
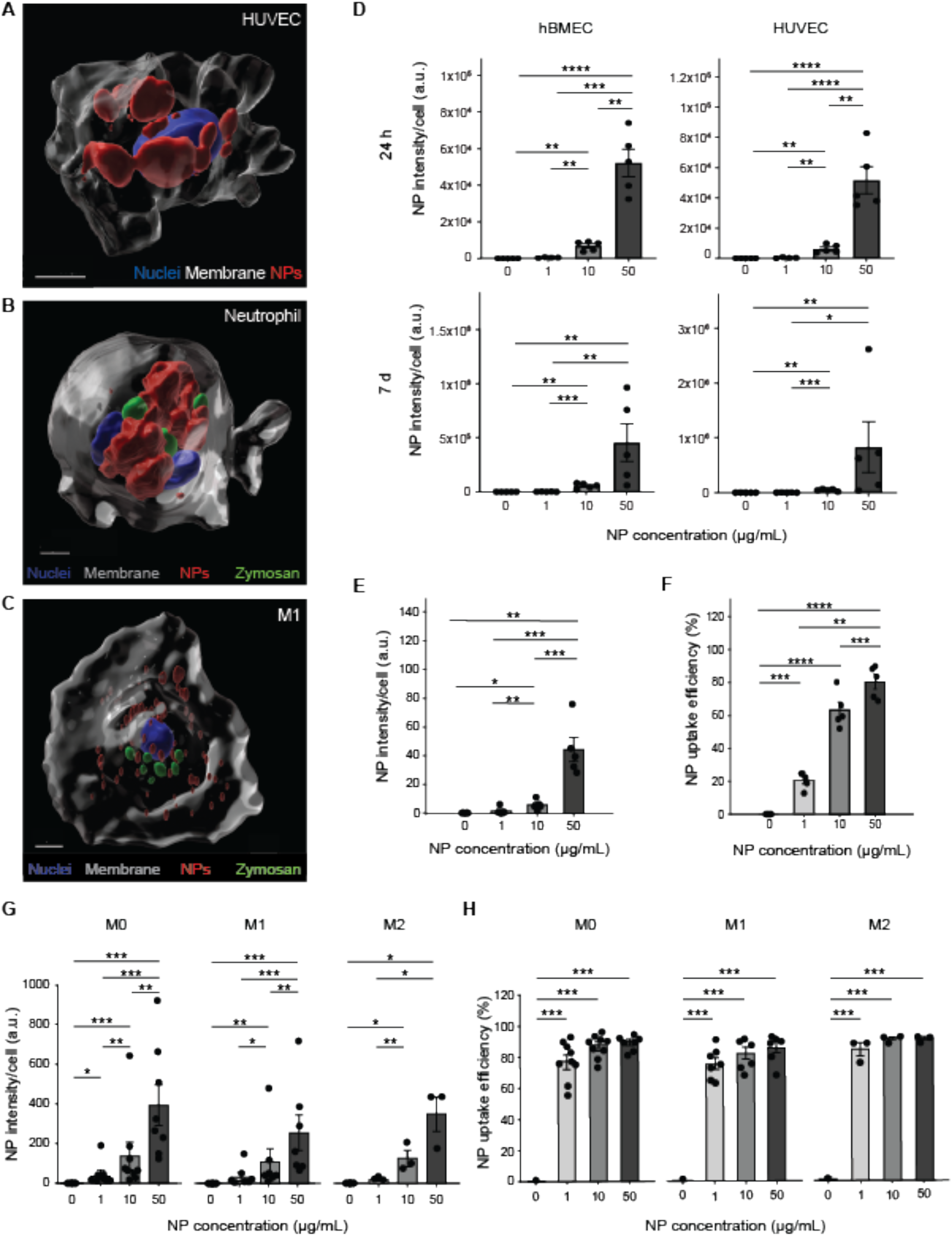
NPs are internalized by endothelial cells, neutrophils, and macrophages in a concentration-dependent manner. **A** Three-dimensional Imaris surface rendering of a HUVEC exposed to 50 µg/mL NPs, showing nuclei (blue), plasma membrane (white) and NPs (red). Scale bar, 15 µm. **B** Three-dimensional rendering of a neutrophil exposed to 50 µg/mL NPs, additionally showing zymosan (green). Scale bar, 4 µm. **C** Three-dimensional rendering of an M1 macrophage exposed to 1 µg/mL NPs. Scale bar, 8 µm. **D** NP fluorescence intensity per cell in hBMECs and HUVECs after 24 h and 7 d of exposure. N = 5, n = 9. **E** NP fluorescence intensity per cell and **F** NP uptake efficiency in neutrophils. N = 5, n = 6. **G** NP fluorescence intensity per cell and **H** NP uptake efficiency in M0, M1 and M2 macrophages. N = 9 (M0; 8 at 50 µg/mL), 7 (M1) and 3 (M2), n = 1–5. NP uptake efficiency is the percentage of cells whose NP fluorescence exceeded the 90th percentile of the matched unexposed cells from the same donor. (**B**, **D**, **E**, **G** and **H**) One-way repeated-measures ANOVA on log2-transformed values with Tukey’s post-hoc comparisons; the plotted bars are background-subtracted, but the statistics were computed before subtraction. Bars show the mean ± s.e.m. with individual biological replicates overlaid. *P < 0.05, **P < 0.01, ***P < 0.001, ****P < 0.0001. N, biological replicates (donors); n, technical replicates (fields of view) per biological replicate and condition.

Uptake in ECs was measured following exposure to NPs alone. In neutrophils and macrophages, NP uptake and phagocytosis were measured simultaneously: cells were exposed to NPs, then to plasma-opsonized zymosan, then fixed. The phagocytosis results from these cells are described in later sections.

In neutrophils and macrophages, internalization was quantified in two ways: NP fluorescence intensity per cell, which reports how much NP each cell contains, and NP uptake efficiency, the percentage of cells whose NP fluorescence exceeded the 90th percentile of the matched unexposed cells from the same donor. In ECs, only NP fluorescence intensity per cell was measured, because discrete borders between cells were difficult to distinguish due to monolayer formation.

In both EC populations, NP fluorescence intensity per cell increased significantly with each increase in concentration after both 1 and 7 days (Fig. 3D). At 50 µg/mL, mean fluorescence after 7 days of exposure was approximately three times higher than after 1 day of exposure. The two timepoints were analyzed separately to preserve donor matching and were not compared statistically, but this difference is consistent with continued accumulation under repeated exposure.

In neutrophils, NP fluorescence intensity per cell increased significantly with each increase in concentration from 1 to 50 µg/mL (Fig. 3E). NP uptake efficiency also increased with concentration, although 10 and 50 µg/mL did not differ from each other (Fig. 3F). Higher NP concentration therefore increased both the number of neutrophils containing NPs and the amount of NP per cell.

In macrophages, NP fluorescence intensity per cell increased significantly with concentration in all three phenotypes (Fig. 3G). Uptake efficiency in macrophages also increased with concentration in all three phenotypes, with 1, 10, and 50 µg/mL all higher than 0 µg/mL but not different from each other (Fig. 3H). Nearly all macrophages had internalized NPs when exposed to 1 µg/mL, so higher concentrations increased the amount of NP per cell rather than the number of cells containing NPs.

### NPs do not impair neutrophil phagocytosis but reduce intracellular MPO at high concentration

Phagocytes have a limited capacity for ingestion; therefore, we hypothesized that cells already carrying high numbers of NPs would ingest fewer particles. We tested this hypothesis in neutrophils by adding plasma-opsonized zymosan following NP exposure (Fig. 4A). Cells exposed to higher NP concentrations had higher internalized NP levels (Fig. 3), so the cells challenged with zymosan carried a dose-dependent NP burden. Zymosan beads were internalized at all NP concentrations (Fig. S4). Neither zymosan beads per cell nor phagocytosis efficiency, the percentage of cells containing at least one zymosan bead, changed with NP concentration (Fig. 4B-C).

**Figure 4.**
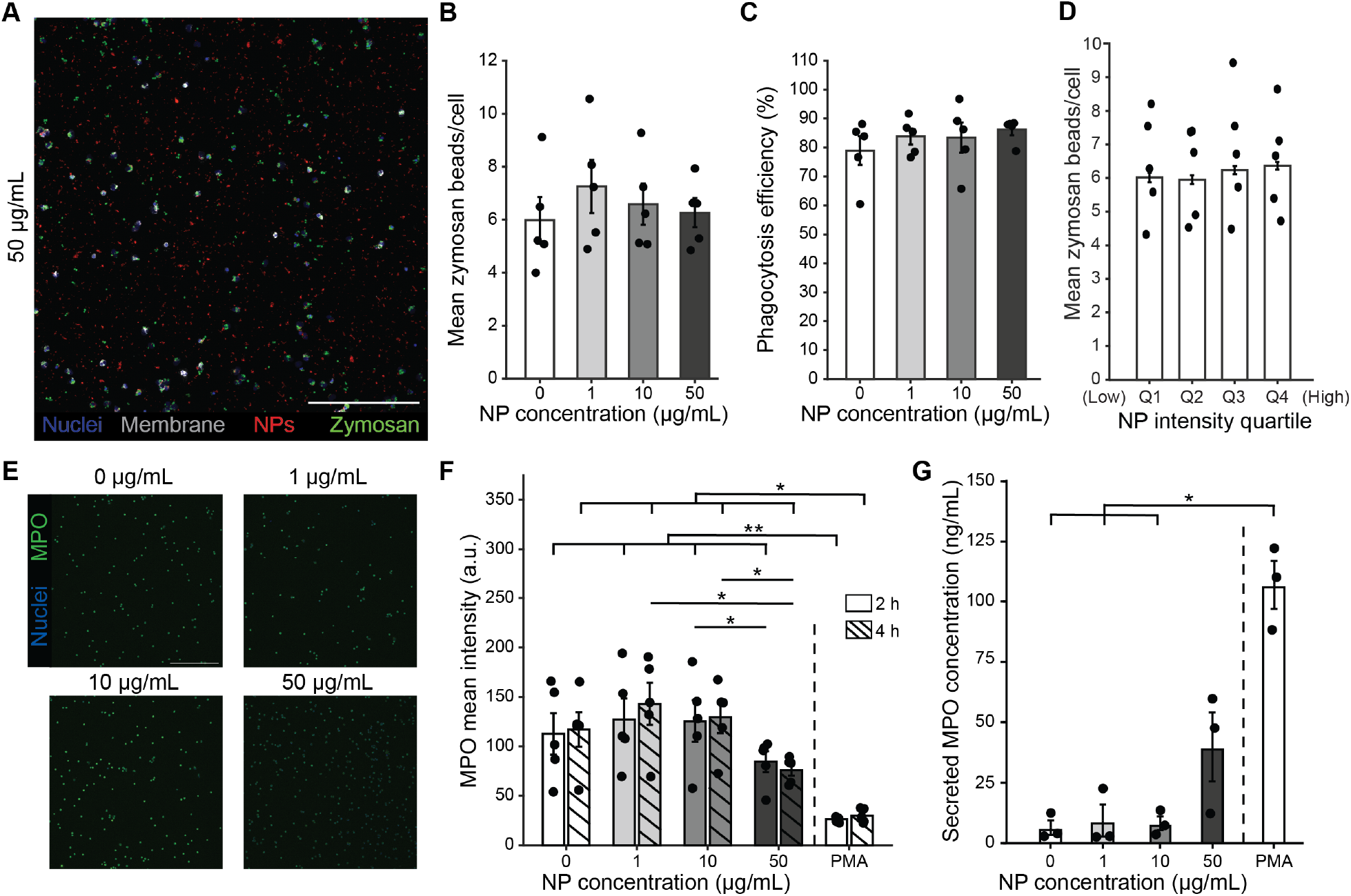
Increasing NP concentration does not affect zymosan phagocytosis but reduces intracellular MPO in neutrophils. **A** Representative image of the NP uptake and phagocytosis assay at 50 µg/mL NPs, showing nuclei (blue), membrane (grey), NPs (red) and zymosan (green). Scale bar, 250 µm. **B** Mean zymosan beads per cell and **C** phagocytosis efficiency, the percentage of cells containing at least one zymosan bead. **D** Mean zymosan beads per cell in quartiles of NP fluorescence intensity (Q1, least NP; Q4, most NP), with quartiles set within each donor across all concentrations. (**B**-**D**) One-way repeated-measures ANOVA on log2-transformed values with Tukey’s post-hoc comparisons. N = 5, n = 6. **E** Representative images of neutrophil MPO after exposure to 0, 1, 10 or 50 µg/mL NPs; nuclei (blue) and MPO (green). Scale bar, 250 µm. **F** Intracellular MPO intensity 2 and 4 h after exposure to NPs or PMA. One-way repeated-measures ANOVA on log2-transformed values with Tukey’s post-hoc comparisons. N = 5, n = 6. **G** Secreted MPO concentration after 4 h of exposure. One-way repeated-measures ANOVA with Tukey’s post-hoc comparisons. N = 3, n = 1. Bars show the mean ± s.e.m. with individual biological replicates overlaid. *P < 0.05, **P < 0.01. N, biological replicates (donors); n, technical replicates (fields of view) per biological replicate and condition.

Population means could conceal an effect confined to the most heavily loaded cells. Therefore, within each donor we ranked cells by NP fluorescence intensity and divided them into quartiles, from the 25% of cells containing the least NP (Q1) to the 25% containing the most (Q4), and took the mean of each quartile as that donor’s value. Because quartiles are set within each donor and pool all concentrations, this analysis asks whether the amount of NP within a cell, rather than the concentration applied, affects phagocytosis. If internalized NPs impaired phagocytosis, cells in Q4 would contain fewer zymosan beads than cells in Q1. Zymosan beads per cell did not differ between quartiles (Fig. 4D).

MPO is the most abundant enzyme in neutrophil granules. It kills ingested microbes but damages host tissue when released, and it contributes to NET formation^24^. To determine whether NPs alter neutrophil MPO, we exposed neutrophils to 0, 1, 10 or 50 µg/mL NPs for 2 or 4 h and measured intracellular MPO by immunofluorescence, with PMA as a positive control for NET induction (Fig. 4E). Neutrophils exposed to 50 µg/mL NPs had lower mean MPO intensity than those exposed to 10 µg/mL at 2 and 4 h, and those exposed to 1 µg/mL at 4 h (Fig. 4F). Lower intracellular MPO could reflect either reduced MPO content or release of MPO from the cell. To distinguish between these possibilities, we measured MPO in the supernatant after 4 h (Fig. 4G). Supernatant MPO increased significantly only in the PMA control relative to 0, 1 and 10 µg/mL; however, secreted MPO following NP exposure showed a positive trend, with higher MPO secretion when exposed to 50 µg/mL. MPO secretion when exposed to 50 µg/mL was also distinctly higher than at lower NP concentrations, although this difference was not significant. Intracellular MPO therefore decreased at the highest NP concentration while supernatant MPO increased, although the latter difference was not significant.

### NP exposure reduces endothelial mediator secretion and neutrophil adhesion to inflamed endothelium

Endothelial cells control leukocyte recruitment through adhesion molecules such as ICAM-1, through chemokines secreted into the vessel lumen, and through VE-cadherin junctions that maintain the barrier between blood and tissue. Inflammatory activation of the endothelium is typically marked by increased ICAM-1 expression and loss of VE-cadherin continuity. HUVECs were chosen as a representative EC model. HBMECs were included as the brain has the highest NP load of any organ ^5,6,25^. To determine whether NP exposure caused endothelial inflammatory activation, we measured ICAM-1 and VE-cadherin by immunofluorescence, quantifying barrier continuity from binary masks of VE-cadherin as the longest continuous junction line per field (Fig. S6A). Neither ICAM-1 expression nor VE-cadherin continuity changed significantly after 1 or 7 days of NP exposure in either cell type (Fig. S6B and C). However, VE-cadherin continuity tended to decrease with increasing NP concentration after 7 days of exposure, more markedly in hBMECs.

We next tested whether NP exposure altered endothelial secretion and subsequent immune cell adhesion. HUVECs were exposed to 0, 10, and 50 µg/mL NPs for 1 day, and to 0, 1, and 10 µg/mL for 7 days. hBMECs were exposed to 0, 10, and 50 µg/mL for 1 day. Conditioned media was collected, and 20 secreted mediators were measured by multiplex immunoassay (MAGPIX). Only MCP-1, IL-8, IL-6, IP-10, IL-1α, and MIP-1β were detectable in all four biological replicates across all conditions. MCP-1, IL-8, and IL-6 directly regulate immune cell recruitment, while IP-10, IL-1α, and MIP-1β are pro-inflammatory mediators that can promote recruitment. Statistical comparisons were made vwithin timepoints only; dose and time were evaluated as contributing factors for HUVECs only.

In HUVECs exposed to NPs for 1 day, MCP-1, IL-8, IL-6, IL1-α, and MIP-1β all decreased with increased NP dose. After 7 days of exposure, IL-6, IL-1α, and MIP-1β also showed decreases in cytokine secretion (Fig. 5A). Exposure time was a significant factor in determining cytokine secretion across all 6 cytokines. Dose was a significant factor for MIP-1β and IL-8. hBMECs also showed decreases in MCP-1, IL-8, IL-6, and IP-10 after 1 day of NP exposure (Fig. 5B).

**Figure 5.**
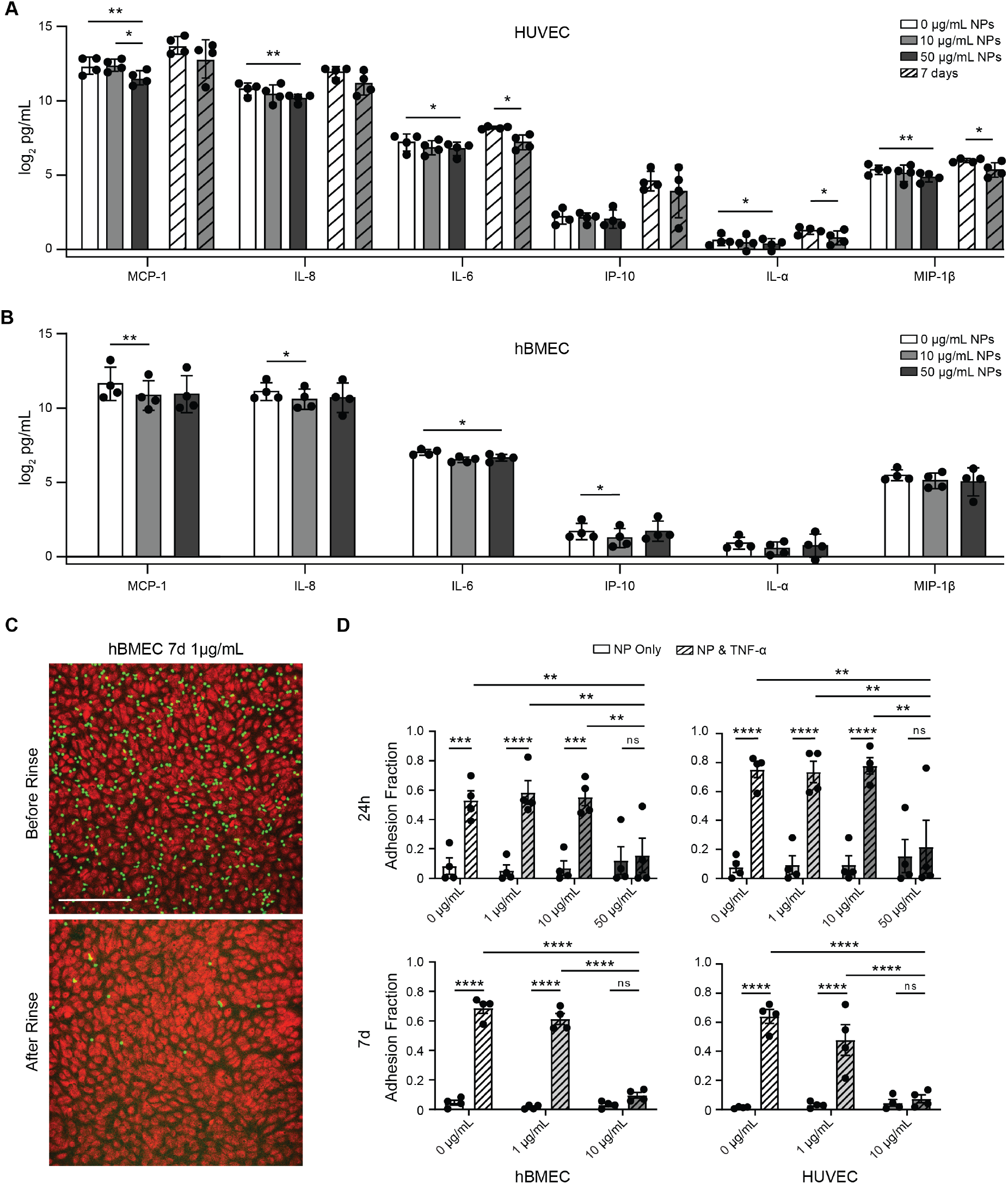
NP exposure reduces endothelial mediator secretion and neutrophil adhesion to inflamed endothelium. **A** Concentrations of 6 analytes in HUVEC conditioned media after 24 h (plain bars) and 7 d (hashed bars) of daily exposure to 0, 10 or 50 µg/mL non-fluorescent NPs, shown as log2 pg/mL. N = 4, n = 1. **B** The same 6 analytes in hBMEC conditioned media after 24 h. N = 4, n = 1. (**A** and **B**) Lognormal one-way ANOVA with Tukey’s post-hoc comparisons; comparisons were made within each timepoint, and HUVEC and hBMEC were not compared with each other. **C** Representative image of neutrophil adhesion before and after rinsing. Scale bar, 250 µm. **D** Adhesion fraction of neutrophils to hBMEC or HUVEC endothelium exposed to NPs for 24 h or daily for 7 d, followed by a secondary TNF-α challenge or media control. Two-way ANOVA with Tukey’s post-hoc comparisons. N = 4, n = 4–5. Bars show the mean ± s.e.m. with individual biological replicates overlaid. ns P > 0.05, *P v< 0.05, **P < 0.01, ***P < 0.001, ****P < 0.0001. N, biological replicates (donors); n, technical replicates (individual wells) per biological replicate and condition.

To determine whether these changes affected the capacity of the endothelium to recruit neutrophils, we measured neutrophil adhesion. Endothelial monolayers were exposed to NPs and stimulated with either 0 or 50 ng/mL TNF-α to model NP only and inflammatory conditions respectively. Adhesion was calculated as the fraction of neutrophils remaining after rinsing (Fig. 5C). NP exposure alone, without inflammatory signaling, did not alter neutrophil adhesion at either time point (Fig. 5D). Secondary activation with TNF-α after 1 day of NP exposure led to increases in neutrophil adhesion in both EC types exposed up to 10 µg/mL, but not when exposed to 50 µg/mL. After 7 days, TNF-α increased adhesion for cells exposed to 0 and 1 µg/mL, but not 10 µg/mL (Fig. 5D). Exposure to 50 µg/mL NPs at 1 day, and to 10 µg/mL at 7 days, reduced neutrophil adhesion compared with lower NP concentrations under TNF-α-stimulated conditions. Collectively, these data suggest that exposure to higher concentrations of NPs limits neutrophil adhesion following inflammation.

The effect of NP exposure on neutrophil adhesion only became apparent under inflammatory conditions. Decreased neutrophil adhesion together with decreased secretion of recruiting mediators indicates that NP exposure reduces the capacity of the endothelium to recruit immune cells to an inflamed site.

### NP exposure does not alter macrophage phagocytosis but increases M1 macrophage cytokine secretion

Macrophages clear pathogens and apoptotic debris in the tissue and regulate inflammation largely through the secretion of cytokines and chemokines. We, therefore, determine how NP exposure altered macrophage phagocytosis and cytokine secretion. As with neutrophils, macrophages were first exposed to NPs and then to plasma-opsonized zymosan. Macrophages followed the same pattern as neutrophils and had increasing uptake efficiency with increasing NP concentration, although the efficiency values were much higher. At 50 µg/mL, intracellular NPs formed large clusters around the nucleus (Fig. 6A and Fig. S4). Three-dimensional renderings of M0 and M2 macrophages showed the same pattern, and zymosan was intracellular at every concentration in all three phenotypes (Fig. S4). Macrophages change shape when activated; therefore, we measured cell area as a marker of macrophage spreading. Cell area did not change significantly with NP concentration in any phenotype (Fig. S7B).

**Figure 6.**
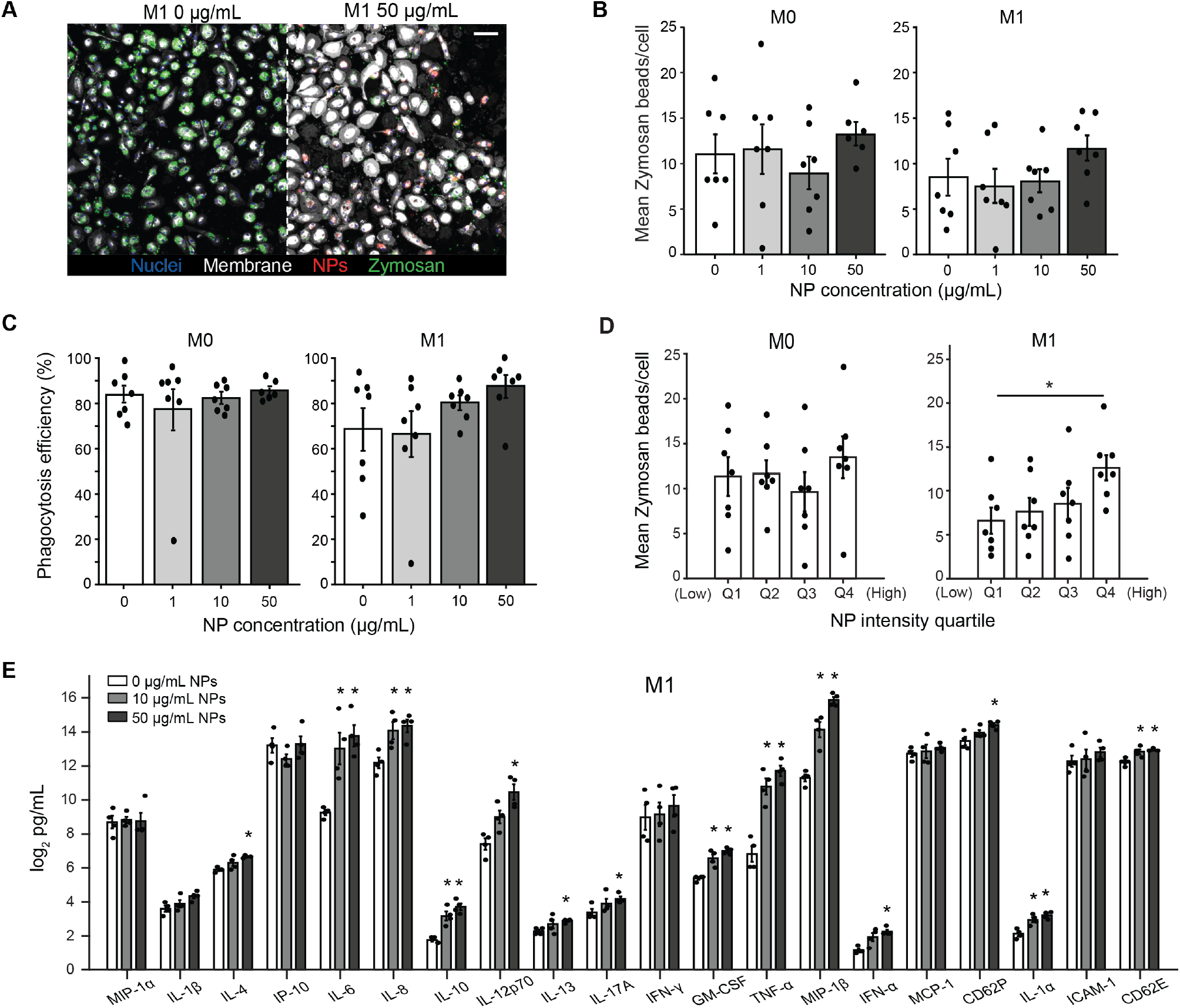
NP exposure does not alter macrophage phagocytosis but increases M1 macrophage cytokine secretion. **A** Representative confocal maximum-intensity projection of M1 macrophages exposed to 0 or 50 µg/mL NPs and then to opsonized zymosan, showing nuclei (blue), plasma membrane (white), NPs (red) and zymosan (green). Scale bar, 30 µm. **B** Zymosan beads per cell and **C** phagocytosis efficiency, the percentage of cells containing at least one zymosan bead, in M0 and M1 macrophages. **D** Mean zymosan beads per cell in quartiles of NP fluorescence intensity (Q1, least NP; Q4, most NP), with quartiles set within each donor across all concentrations. (**B** to **D**) One-way repeated-measures ANOVA on log2-transformed values with Tukey’s post-hoc comparisons. N = 7 (M0; 6 at 50 µg/mL, and M1), n = 1–5. **E** Concentrations of all 20 analytes in conditioned media of M1 macrophages after 24 h exposure to 0, 10 or 50 µg/mL non-fluorescent NPs, quantified by multiplex immunoassay (MAGPIX, 20-plex) and shown as log2 pg/mL. Each concentration was compared with the matched 0 µg/mL well from the same donor by two-sided paired t-test on log2-transformed concentrations, with Benjamini-Hochberg correction across all 20 analytes and both concentrations. N = 4, n = 1. Bars show the mean ± s.e.m. with individual biological replicates overlaid (**B** to **D**) or the mean ± s.e.m. of log2 concentrations, which is the geometric mean (**E**). In **B** to **D**, *P < 0.05; in **E**, *adjusted P < 0.05. N, biological replicates (donors); n, technical replicates per biological replicate and condition.

Phagocytes have a limited capacity for ingestion, so we next asked whether macrophages already carrying NPs would ingest fewer particles. Neither zymosan beads per cell nor phagocytosis efficiency changed with NP concentration in M0 or M1 macrophages (Fig. 6B-C). M2 macrophages were not included, because the clearance function that defines the alternatively activated state is efferocytosis rather than the ingestion of exogenous particles^8^. Like neutrophils, cells were ranked by NP fluorescence intensity and divided into quartiles within each donor, and the mean number of zymosan beads per cell was calculated for each quartile and donor. In M1 macrophages, the cells in Q4 contained more zymosan beads than those in Q1. In M0 macrophages, bead counts did not differ between quartiles (Fig. 6D). Internalized NPs therefore did not reduce phagocytosis even in the most heavily loaded macrophages.

Macrophages regulate the inflammatory environment through cytokine secretion, and their secretion profile depends on polarization state^21,22^. Therefore, we measured the secretory profile of M0, M1, and M2 macrophages following NP exposure. Macrophages were exposed to 0, 10, or 50 µg/mL NPs for 24 h, and the concentration of 20 cytokines, chemokines and soluble adhesion molecules in the conditioned media was measured by multiplex immunoassay (MAGPIX). Baseline secretion differs between phenotypes, so each phenotype was compared to the donor-matched 0 µg/mL well. All 20 analytes are shown for M1 macrophages in Fig. 6E and for M0, M1 and M2 macrophages in Fig. S8.

M1 macrophages responded broadly to NPs: 14 of the 20 analytes increased significantly following NP exposure, eight cytokines increased at both 10 and 50 µg/mL of NP exposure (Fig. 6E). The largest increases were in classical inflammatory mediators. TNF-α, MIP-1β, and IL-6. IL-8, IL-1α, GM-CSF, and soluble E-selectin (CD62E) increased following exposure to both 10 and 50 µg/mL of NPs. IL-12p70, IFN-α, IL-17A, soluble P-selectin (CD62P), IL-4, and IL-13 increased following exposure to 50 µg/mL only. The response was not restricted to pro-inflammatory mediators: the anti-inflammatory cytokine IL-10 also increased at both concentrations. IL-1β, MIP-1α, MCP-1, IP-10, IFN-γ, and soluble ICAM-1 did not change significantly.

Neither M0 or M2 macrophages showed significant alterations in secretion with NP exposure to 10 or 50 µg/mL NPs (Fig. S8A and C). Several analytes, including TNF-α, GM-CSF, IL-13, and IFN-α, were at or near the lower limit of quantification in some or all M0 and M2 donors, so small changes in these analytes could not be detected.

All three phenotypes internalized NPs in a concentration-dependent manner, but only M1 macrophages, which were activated with LPS and IFN-γ before exposure, altered their secretion. The cytokine response to NPs therefore changed with the activation state of the cell.

## Discussion

The first step in understanding whether NP exposure contributes to the onset of autoimmunity is determining its effect on innate immune activation^3^. Here, we measured functional responses of primary human neutrophils, macrophages, and endothelial cells following exposure to physiologically relevant concentrations^4^. We found all cell types internalized NPs in a concentration-dependent manner without a loss in viability (Fig. 2-3), yet the functional responses diverged. Neutrophils contained less intracellular MPO following exposure to the highest NP concentration (Fig. 4F-G). M1 macrophages increased secretion of 14 of 20 inflammatory mediators while M0 and M2 macrophages did not (Fig. 6E and Fig. S8). ECs did not show signs of inflammation (Fig. S6) but instead had reduced secretion of leukocyte-attracting chemokines and supported less neutrophil adhesion once inflamed (Fig. 5). Therefore, NP exposure alone did not cause a universal pro-inflammatory response, instead the type and degree of dysregulation differed by cell population. Further, each of the three responses we measured can be connected to immune dysregulation implicated in autoimmune disease.

Our results contradict the majority of existing literature which report high inflammatory responses to NP exposure regardless of cell type or activation. Studies often miss critical aspects of the immune responses by not investigating multiple cell types at once^12^, using bare particles rather than biocorona-coated NPs as would be seen *in vivo*^13,14^, and using supraphysiological concentrations tied to cell death^4,26^. Under these conditions, the reported response is consistently inflammatory; however, it is typically accompanied by reduced viability^9,11,27^ so reported inflammatory responses are likely impacted by cell death. Our study design eliminates these issues; using multiple primary human cells, identical biocorona-coated NPs, and physiologically relevant concentrations of NPs we see distinct inflammatory and functional responses. Importantly, we see high viability across all cell types meaning the responses we observe can be attributed to cell identity and activation state rather than cell death. It also allowed us to extend endothelial exposures to 7 days, longer than has been reported previously *in vitro*, and to work with neutrophils, which are already prone to apoptosis^19^.

MPO is the most abundant neutrophil granule protein and damages host cells and tissue when released, stimulating neutrophil recruitment and increased inflammation^20,24,28^ It is also the target autoantigen in one of the two forms of anti-neutrophil cytoplasmic antibody (ANCA)-associated vasculitis^29^ and only becomes available as an autoantigen after it is released by the cell^28^. Increased concentrations of plasma MPO have also been reported in inflammatory autoimmune diseases including rheumatoid arthritis and multiple sclerosis ^29–32^. To our knowledge, this is the first study to measure MPO release in primary human neutrophils following exposure to NPs at concentrations reported in human blood^33^.

Neutrophils exposed to the highest NP concentration contained less intracellular MPO (Fig. 4F), and supernatant MPO was higher at the same NP exposure concentration, although this difference was not significant (Fig. 4G). Together, these results suggest neutrophils release MPO following exposure to high NP concentrations. The increased availability of MPO matches autoimmune states seen across various diseases^28–31^. Our results suggest that NP-driven MPO release by neutrophils is one plausible mechanism by which NP exposure could contribute to autoimmune states.

Macrophages dictate the inflammatory state of a tissue through cytokine secretion^21,22^. Persistent M1-like macrophages are a feature of systemic lupus erythematosus, where they sustain inflammation that drives tissue damage^34,35^. Furthermore, M1 macrophage cytokine secretion is directly linked to chronic inflammation and generalized autoimmune disease: TNF-α, IL-6 and GM-CSF are all therapeutic targets in inflammatory rheumatic disease^36–38^, and IL-1α, IL-8 and MIP-1β sustain the recruitment of further leukocytes into the tissue^18^. It is, therefore, most important to understand the M1 response to NPs in the context of autoimmune disease. The literature reports that all macrophages show increased inflammation following NP exposure regardless of phenotype^9,10,39^. However, M0, M1, and M2 macrophages internalize nanoparticles at different rates and the polarization state of a macrophage impacts its response^21,22,40,41^; therefore, we expected the response to NPs to differ by phenotype. In fact, we only observed an increase in inflammatory cytokines in M1 macrophages, in which 14 of the 20 analytes measured increased (Fig. 6E). The largest increases, relative to the 0 µg/mL control, were in TNF-α, MIP-1β, IL-8 and IL-6. All aforementioned cytokines along with IL-1α, GM-CSF, IL-10, and soluble E-selectin increased following exposure to both 10 and 50 µg/mL of NPs. All cytokines with increased secretion are established drivers of autoimmune inflammation. Overall, NP exposure increased inflammatory secretion in M1 macrophages, the predominant polarization state in inflamed tissue. Therefore, we expect increases in inflammation to be most pronounced in the already inflamed tissue of autoimmune disease, although whether it occurs *in vivo* remains to be tested.

Endothelial cells regulate the immune response by mediating leukocyte migration into tissues through chemokine secretion and adhesion molecule expression, such as ICAM-1 ^17,18^. In systemic lupus erythematosus and rheumatoid arthritis that control is perturbed: the endothelium is injured and activated, barrier integrity is compromised and immune cell recruitment increases^42^. Current literature examining the effects of NPs on ECs aligns with autoimmune dysfunction in that NP exposure leads to barrier disruption^27,43,44^. However, the degree to which the barrier is disrupted, the timescale it happens on, and the cell death associated greatly varies with the NP conditions used. Collectively, the literature emphasizes the importance of NP type on the EC response^45,46^. Our choice in NP and resulting high viability may explain why we see no EC barrier disruption as measured by ICAM-1 expression and VECAD continuity (Fig S6). However, our work highlights another important variable in addition to NP type: inflammatory state, especially when evaluating endothelial-immune cell interactions. We found decreases in neutrophil adhesion following NP exposure to the highest concentrations required a secondary TNF-α challenge (Fig 5D). Like macrophages, inflammation can impact cell response to NP exposure. Further, to our knowledge, this is the first study to investigate how a secondary inflammatory stimulus impacts immune cell recruitment to an endothelium, and one of few to look at endothelial-leukocyte interactions upon NP exposure^43,47,48^. Endothelial leukocyte interactions upon NP exposure are understudied, but to have a complete picture of NP impact, relevant inflammatory activation will be critical along with NP choice.

While we addressed several challenges in understanding how NPs affect immune cells, several limiting factors are important to consider. In this study, we only used pristine NPs, not weathered NPs. NPs in the environment are weathered by ultraviolet light and mechanical abrasion, which changes particle shape, oxidizes the particle surface, and increases cytotoxicity relative to pristine particles^26,49,50^ Therefore, the responses reported here are likely less inflammatory than weathered NPs. Additionally, all of the NPs used were uniform in size. Environmental nanoplastics span a broad size distribution, and size determines both the route of internalization and the number of particles a cell takes up at a given mass concentration^9,26^. Consequently, a monodisperse exposure cannot reproduce the internalized dose a polydisperse exposure would deliver. We also only studied polystyrene NPs, not the range of plastic types that would be found in the body. Polymer chemistry determines surface charge, the proteins recruited into the biocorona, and the additives carried with the particle which can affect cell viability^13,51^, so these results describe polystyrene of approximately 0.5 µm in diameter rather than nanoplastics as a collective. A further constraint is that in some neutrophil and macrophage experiments we measured the response to NP exposure alone, without a subsequent inflammatory stimulus like TNF-α. The endothelial experiments show that the functional consequences of NP exposure can appear only after such a stimulus, so the phagocyte responses reported here may underestimate what occurs in an inflamed tissue. Finally, our primary cells came from donors of both biological sexes, but the data were not analyzed by sex. Four of every five people that develop an autoimmune disease are women and over 100 different autoimmune diseases disproportionately affect women^1,52,53^. Thus, whether the innate immune response is dysregulated by NPs could depend on the biological sex of the cells. Lastly, studying these cells in additional inflammatory contexts would also distinguish whether low NP doses prime cells towards an inflammatory phenotype, indicating a predisposition towards autoimmunity, or act only to exacerbate established disease.

In summary, we sought to determine how exposure to physiologically relevant NP concentrations affected innate immune-related cell function. Neutrophils showed evidence of MPO release, only M1 macrophages increased secretion of cytokines known to drive autoimmune disease, and endothelial cells only exhibited decreased neutrophil adhesion with prior exposure to TNF-α. Therefore, the functional consequence of NP exposure depends on the exposed cell population and on the inflammatory environment surrounding the cell. Determining how innate immune-related cell function is altered by NP exposure is the first step in establishing a connection to autoimmune disease. While further investigation is required to define the connection between NP exposure and autoimmune disease, our results demonstrate that NPs can dysregulate innate immune-related cell function.

## Methods

### Study approval

All blood samples were collected from willing donors aged 18 to 65 under a protocol approved by the University of Colorado Boulder Institutional Review Board (#20-0082), in accordance with the Declaration of Helsinki. Written informed consent was obtained from every donor. Blood was collected into BD Vacutainer EDTA tubes (366643, BD Biosciences). Donors were not selected or stratified by sex or age beyond the eligibility range above.

### Biocorona-coated nanoplastics preparation

Non-fluorescent nanoplastics with a biocorona were prepared by adding 10 µL of polystyrene beads (LB5, Sigma-Aldrich) to 1 mL of human plasma collected as a byproduct from human blood draws, mixing well with a pipette, and then incubating at 37 °C for 24 h before use. For use in experiments, nanoplastics were centrifuged at 17,000 × g for 15 min and were resuspended in either EGM-2 medium (for neutrophil and endothelial cell experiments) or AIM-V (for macrophage experiments) and diluted to the desired concentration. Fluorescent nanoplastics with a biocorona were prepared by the same method, except using 10 µL of carboxylate-modified fluorescent red polystyrene beads (L3280, Sigma-Aldrich) in place of the non-fluorescent beads.

Both preparations used pristine and unweathered nanoplastics, supplied as aqueous suspensions and not subjected to artificial aging, UV, or mechanical weathering. The fluorescent red carboxylate-modified beads (L3280) have a reported mean diameter of 0.5 µm (λex ≈ 575 nm, λem ≈ 610 nm, imaged with 561 nm excitation) and were used for all NP uptake experiments, whereas the unlabeled beads (LB5) have a reported mean diameter of 0.46 µm and were used for the conditioned-media experiments, because fluorescence from the labelled particles would otherwise interfere with the bead-based immunoassay readout. Stocks were vortexed and bath-sonicated immediately before use to disperse aggregates.

### TEM/SEM image methods

Biocorona-coated nanoplastic samples that were incubated for 24 h and resuspended in deionized water were negatively stained by the drop method in the following manner: 5 µL aliquots of sample and 2% uranyl acetate aqueous stain applied sequentially to a freshly glow-discharged carbon-coated 200-mesh copper grid. Images were collected on an FEI Tecnai T12 Spirit operating at 100 kV with a LaB6 filament and equipped with an AMT 2k x 2k side-mounted CCD camera.

### Hydrodynamic diameter and zeta potential

Non-fluorescent nanoplastics (30 µL) or fluorescent nanoplastics (30 µL) were incubated in 1 mL of human plasma for 10 min or 24 h to form a biocorona, or left uncoated as a control. NPs were then centrifuged at 17,000 × g for 20 min, the plasma was aspirated, and the NPs were resuspended in 2.5 mL of water. The suspension was transferred to an Omega cuvette (225288, Anton Paar) and the zeta potential was measured on an Anton Paar LiteSizer 500 with an equilibration time of 2 min, a target temperature of 25 °C, and water as the solvent. The Omega cuvette was washed thoroughly (three times with deionized water, then once with methanol, and completely dried using an air blow gun) between measurements. Each suspension was then diluted into a 1.5 mL cuvette (14955127, Fisher) and the hydrodynamic diameter was measured on the same instrument with a 2 min equilibration time, a target temperature of 25 °C, the material set to polystyrene latex, and water as the solvent. Plasma from a different donor was used for each biological replicate.

### Endothelial cell culture

Pooled human umbilical vein endothelial cells (HUVEC, 50-305-964, PromoCell) were cultured using Endothelial Growth Medium (EGM-2, NC9525043, Lonza Walkersville CC3162) at 37 °C and 5% CO₂. Cells were expanded in T-75 flasks and passaged at 80% confluency. HUVECs were used between passages 4 to 7.

Human brain microvascular endothelial cells (hBMEC, ACBRI 376, Cell Systems) were cultured using EGM-2 at 37 °C and 5% CO₂. Cells were cultured on top of fibronectin coated (2µg/cm^2^, 8248 ScienCell) TC plastic. Cells were expanded in T-75 flasks and passaged at 80% confluency. hBMECs were used from passages 5 to 8.

### Isolation of primary human leukocytes

Primary human neutrophils were isolated from whole blood samples with the MACSxpress Neutrophil Isolation Kit (130-104-434, Miltenyi Biotec) followed by the Erythrocyte Depletion Kit (130-098-196, Miltenyi Biotec) according to the manufacturer’s protocols.

Peripheral blood mononuclear cells were isolated by density-gradient centrifugation. Whole blood was diluted 1:1 in sterile 1× PBS and layered over 15 mL of Ficoll-Paque PLUS (Cytiva, 17144002) in 50 mL conical tubes, then centrifuged at 400 × g for 35 min at room temperature with acceleration and brake set to 1 and 0 to preserve the interface. The buffy coat was collected, brought to 40 mL in PBS, and washed twice at 400 × g for 10 min followed by two spins at 100 × g for 10 min to deplete platelets. Monocytes were purified by negative selection with the Pan Monocyte Isolation Kit (Miltenyi Biotec, 130-096-537) and resuspended in pre-warmed AIM-V medium (Life Technologies, 12055091) at 1.5 × 10⁶ cells/mL before plating.

### Primary macrophage differentiation and polarization

Monocytes were differentiated into macrophages over 6 days in AIM-V (12055091, Thermo Fisher) supplemented with M-CSF (2 ng/mL; 300-25, PeproTech), with medium refreshed every 24 h. Polarisation was induced on day 6 by adding, on top of the M-CSF baseline, either IFN-γ (20 ng/mL; BioLegend) with LPS (100 ng/mL; Sigma, L4516) for M1 or IL-4 (20 ng/mL; 130-093-915, Miltenyi Biotec) for M2; M0 wells received M-CSF alone. Cells were held under these conditions for 24 h before nanoplastic exposure.

### Nanoplastic exposure

NPs were applied at final concentrations of 0, 1, 10 and 50 µg/mL for uptake, viability, neutrophil adhesion, protein expression, and myeloperoxidase experiments, and at 0, 10 and 50 µg/mL for conditioned-medium experiments; control wells received cell medium alone. Bioreplicates are defined as individual donors or distinct EC passages. Within each experiment, doses were applied to cells from the same bioreplicate, so dose comparisons are all paired within bioreplicates.

Macrophages were exposed to NPs carrying an autologous biocorona: 10 µL of the particle stock was diluted in 1 mL of plasma from the same donor whose monocytes were used in that experiment, incubated for 24 h at 37 °C and 5% CO₂, then diluted in AIM-V to the working concentrations. Before dosing, day-6 polarizing medium was aspirated and wells were rinsed with warm AIM-V to clear residual cytokine, after which the corresponding NP suspension was added to each M0, M1 or M2 well and left for 24 h at 37 °C and 5% CO₂.

Neutrophils were resuspended in EGM-2 at 8 × 10⁵ cells/mL and 50 µL was added per well, followed by 50 µL of NP suspension, giving the final concentrations above. Where a positive control was included, 50 µL of 44 µM phorbol 12-myristate 13-acetate (PMA) was added instead of NP suspension.

HUVECs and hBMECs were seeded at 15k cells per well in 48 well plates, and 10k cells per well in 96 wells. 300 µL of NP suspension were added to each well for 96 well plates. 100 µL of NP suspension were added to each well for 96 well plates. Endothelial cells were grown to confluency prior to exposure to NPs. Cells were exposed daily to either 0, 1, 10, and 50 µg/mL of NPs for up to 7 days.

### Neutrophil viability

Neutrophils were resuspended in 1× PBS without calcium or magnesium (10010-049, Thermo Fisher) and stained with Hoechst 34580 (H21486, Thermo Fisher, 1:100) and calcein AM (C3100MP, Invitrogen, 5 µM) for 10 min at room temperature. Cells were centrifuged at 200 × g for 5 min, the stain solution was aspirated, and cells were resuspended in 1 mL of EGM-2 containing propidium iodide (53-706-05ML, Millipore Sigma, 2 µg/mL) and incubated for 15 min at room temperature. Neutrophils were diluted to 8 × 10⁵ cells/mL, plated at 50 µL per well and dosed as described above. Wells were imaged every 30 min for 8 h at 37 °C and 5% CO₂. Live cells were gated as Hoechst-positive and calcein AM-positive only, and dead cells as Hoechst-positive and propidium iodide-positive only. Viability was calculated per field as the number of live cells divided by the total number of live and dead cells.

### Macrophage viability

After 24 h of NP exposure macrophages were stained with Hoechst 34580 (H21486, Thermo Fisher, 1:100) and calcein AM (C3100MP, Invitrogen, 5 µM) in AIM-V for 10 min at room temperature. The stain solution was replaced with AIM-V containing propidium iodide (53-706-05ML, Millipore Sigma, 2 µg/mL) and cells were incubated for 15 min at room temperature. Wells were imaged without fixation at 10x magnification; three wells per condition. Live cells were identified as Hoechst-positive and calcein-positive and dead cells as Hoechst-positive and propidium iodide-positive, and both were counted in NIS-Elements. Viability was calculated per well as live cells divided by the total of live and dead cells, and wells were averaged to one value per donor and condition.

### Endothelial cell viability

Viability was evaluated in 96 well plates. Cell media was removed and cells were stained using Hoechst 34580 (H21486, Thermo Fisher, 1:100), calcein AM (C3100MP, Invitrogen, 5 µM), and propidium iodide (PI, 53-706-05ML, Sigma, 1:500) in EGM2 for 30 min t 37 °C and 5% CO₂. Cells were rinsed 3 times with 1× PBS without calcium or magnesium (10010-049, Thermo Fisher). Wells were imaged at 10x magnification with 3 images per well and 3 wells per condition. Live cells were identified as Hoechst and calcein-positive, and PI negative. Dead cells were any cells staining positive for Hoechst and PI, regardless of calcein-AM presence. Viability was calculated as lives cells divided by the combination of live and dead cells. Thresholds for calcein and PI were set manually for each donor and day and analyzed using NIS elements using a custom macro.

### Nanoplastic uptake

Internalization of fluorescent NPs was assessed in endothelial cells, neutrophils, and M0, M1 and M2 macrophages. Nuclei and plasma membranes were labeled in each cell type before fixation so that NP uptake could be quantified per cell.

Neutrophils were dosed as described above. After 2 h at 37 °C and 5% CO₂, cells were stained with Hoechst 34580 (H21486, Thermo Fisher, 1:100) and Vybrant DiD Cell-Labeling Solution (V22887, Invitrogen, 1:200) for 20 min at 37 °C and rinsed three times with warm 1× PBS.

HUVEC and hBMEC monolayers were grown in 48 well plates and labeled with Vybrant DiD Cell-Labeling Solution (V22887, Invitrogen, 1:200) and Hoechst 34580 (H21486, Thermo Fisher, 1:100) for 20 min at 37 °C and rinsed three times with warm 1× PBS.

Macrophages (1 × 10⁶ cells/mL) were labeled after NP exposure with Vybrant DiD Cell-Labeling Solution (V22887, Invitrogen, 1:1000) and Hoechst 34580 (H21486, Thermo Fisher Scientific, 1:1000) for 20 min at 37 °C and rinsed three times with warm 1× PBS to remove unbound dye.

All wells were fixed in 4% paraformaldehyde (AC416780250, Fisher Scientific) for 15 min at room temperature and washed three times in cold 1× PBS.

### Zymosan phagocytosis

Phagocytic capacity was measured in neutrophils and in M0 and M1 macrophages using the same opsonized zymosan assay. M2 macrophages were excluded from this comparison by design, because the clearance function that defines the alternatively activated state is efferocytosis rather than ingestion of exogenous particles^8^, so a zymosan uptake index would report a function that is not their principal role. Alexa Fluor 488-conjugated Zymosan A BioParticles (Z23373, Thermo Fisher Scientific) were opsonized in normal plasma from the same donor used for the corresponding NP incubation, diluted 1:1 in 1× PBS for 1 h at 37 °C, pelleted at 1,500 × g for 5 min, washed once in EGM-2 or AIM-V to remove residual plasma, and resuspended in EGM-2 or AIM-V. Opsonized zymosan was added after NP exposure and labeling at a particle-to-cell ratio of 4:1 for neutrophils and 10:1 for macrophages. Plates were centrifuged at 100 × g for 1 min to bring particles into simultaneous contact with all cells and incubated for 1 h at 37 °C and 5% CO₂. Non-internalized particles were removed by three washes in cold 1× PBS. Cells were fixed in 4% paraformaldehyde (AC416780250, Fisher Scientific) for 15 min at room temperature, washed three times in cold 1× PBS and stored at 4 °C until imaged.

### Neutrophil myeloperoxidase imaging

Neutrophils were resuspended in EGM-2 at 8 × 10⁵ cells/mL, plated at 50 µL per well and dosed with non-fluorescent NPs or PMA as described above. After 2 h or 4 h at 37 °C and 5% CO₂, 100 µL of 4% paraformaldehyde in 1× PBS was added per well and incubated for 15 min at room temperature, and cells were washed once in 1× PBS. Cells were stained with Hoechst 34580 (H21486, Thermo Fisher, 1:100) for 10 min at room temperature, washed three times in 1× PBS, permeabilized in 150 µL of 0.1% Triton X-100 for 10 min and washed once in 1× PBS. Cells were blocked in 150 µL of 5% BSA in PBS for 1 h and stained for myeloperoxidase (MPO monoclonal antibody, FITC, 11-1299-42, Fisher, 1:1000) at room temperature for 1 h. Cells were washed three times in 1× PBS and stored at 4 °C until imaged. Nuclei were counted in NIS-Elements. MPO masks were generated with a custom MATLAB script (MathWorks) that measured MPO pixel intensity and MPO-positive area.

### Neutrophil myeloperoxidase secretion

Neutrophils were resuspended in EGM-2 at 8 × 10⁵ cells/mL, plated at 50 µL per well and dosed with non-fluorescent NPs or PMA as described above. After 4 h at 37 °C and 5% CO₂, the supernatant was collected, centrifuged at 17,000 × g for 5 min to remove NPs and cells, and frozen at −20 °C until assayed. Secreted MPO was quantified with a Quantikine ELISA kit (DMYE00B, R&D Systems) on supernatants diluted tenfold, according to the manufacturer’s protocol.

### Endothelial cell Alamar blue

Endothelial cells were plated in 96 well plates. ECs were treated with Alamar blue reagent (502521743, Fisher) per manufacturer instructions and incubated at 37 °C and 5% CO₂ for 2 hours before fluorescent readout was measured on a fluorescent plate reader. Blanks wells acted as controls for background fluorescence and were subtracted from all other readouts.

### Endothelial staining of ICAM-1, VE-cadherin, F-actin, cell count, and coverage

Endothelial cells were plated in 48 well plates. ECs were fixed in 2% PFA for 15 min at room temperature and washed 3 times with 1% BSA in 1× PBS without calcium or magnesium. Cells were permeabilized with 0.1% Tween 20 for 10 minutes at room temperature and washed 3 times with 1% BSA in 1× PBS without calcium or magnesium. Fixed and permeabilized cells were stained overnight at 4 °C with Hoechst (1:200), Phalloidin (1:1000, 501964078, Fisher), anti-VECAD (5:600, 130-125-985, Miltenyi Biotech), and anti-ICAM (1:200, BBA20, R&D Systems) in 1% BSA solution. Staining solution was aspirated the next morning, and cells were washed 3 times with 1× PBS and stored up to 3 days at 4 °C before imaging. Cells were imaged using a 20x WI objective with 3 wells per condition.

F-actin coverage of the field of view was used to determine cell spreading. A mask of f-actin coverage was made using Otsu thresholding. The resulting mask was used to determine where ICAM presence would be quantified. A skeleton was made of VE-cadherin staining, and the longest line recorded in the resulting skeleton was used as a proxy for endothelial barrier continuity. Number of nuclei were quantified using bright spots counting of Hoechst. All image processing was done in NIS elements using a custom macro.

### Neutrophil static adhesion to endothelium

Endothelial cells were plated in a 96 well plate and were exposed to either 0 or 50 ng/mL of TNF-α for 2 hours. Isolated neutrophils were stained with calcein-AM (5 µM) for 10 min at room temperature. 3 × 10^6^ neutrophils were added to each well for 15 min. Neutrophils were rinsed using media 3 times. Images were taken after neutrophil loading and after rinsing at 10x objective. Adhesion fraction was defined as the number of neutrophils present before rinsing over the number of neutrophils present after rinsing. Neutrophil cell count was quantified using a custom MATLAB script.

### Multiplex cytokine quantification in conditioned medium

Supernatants were collected 24 h after exposure to unlabeled NPs (0, 10 and 50 µg/mL) from M0, M1 and M2 macrophage wells and from HUVEC and hBMEC monolayers, cleared of debris at 400 × g for 10 min and stored at −80 °C until analysis. Samples were diluted 1:2 in the culture medium appropriate to the cell type of origin, AIM-V for macrophages and EGM-2 for endothelial cells.

Within the available assay capacity, conditions were prioritized to allow a 1 versus 7-day comparison at matched doses in HUVEC and a 1-day comparison between the two endothelial types; hBMEC 7-day conditions and all 1 µg/mL doses were therefore not included. Similarly, macrophages at 1 µg/mL doses were also not included.

Cytokines, chemokines and soluble adhesion molecules were quantified with the Human ProcartaPlex Inflammation Panel 20-plex (Thermo Fisher Scientific) on a Luminex MAGPIX according to the kit protocol, against a seven-point standard curve prepared by 1:4 serial dilution of the pooled standards. Standard curves were fitted per analyte with a five-parameter logistic model and concentrations were back-calculated with the twofold dilution factor applied. Duplicates with a coefficient of variation above 15%, or above 20% near the lower limit of quantification, were excluded, and values below the lowest standard were assigned that analyte’s lower limit of quantification. A medium-only blank was run for each culture medium and subtracted from the corresponding samples.

### Confocal image acquisition

Confocal images were acquired on a Nikon A1R HD25 laser scanning confocal microscope mounted on a Nikon Ti2-E inverted stand in galvano scanning mode, using Nikon Elements acquisition software. Neutrophil, macrophage, and endothelial cells viability were imaged live with a Nikon 10×/0.45 NA air objective. Endothelial cells and neutrophils were maintained at 37 °C, 5.0% CO₂ and 90% relative humidity in an OKO Labs environmental chamber during time-lapse imaging. All other experiments were imaged after fixation. The neutrophil MPO experiment was imaged with a Nikon Apo LWD 20×/0.95 NA water-immersion λS objective.

The NP uptake and zymosan phagocytosis experiments were imaged in four channels per field, with sequential excitation at 405 nm for Hoechst 34580 (nuclei), 488 nm for Alexa Fluor 488-zymosan, 561 nm for the fluorescent NPs and 640 nm for DiD (plasma membrane). Fields were acquired with either a Nikon 20×/0.75 NA air objective or a Nikon Apo LWD 20×/0.95 NA water-immersion λS objective across experimental replicates, and masking parameters were calibrated to the corresponding pixel size as described below. Confocal z-stacks for three-dimensional reconstruction were acquired on the same instrument with the Nikon Apo LWD 20×/0.95 NA water-immersion λS objective at a z-step of 0.1 µm, which oversamples relative to the axial resolution of the objective and was chosen to support accurate surface reconstruction.

### Three-dimensional reconstruction and confirmation of nanoplastic internalization

Surface-adherent particles cannot be distinguished from internalized particles in a single optical plane, so NP internalization was verified in three dimensions in all cell types. Z-stacks spanning the full cell depth were acquired as described in the preceding section, imported into Imaris (Oxford Instruments, version 11.0.1) and rendered in three dimensions. All four channels were reconstructed as surfaces rather than as spots, so that the plasma membrane (DiD, 640 nm), nucleus (Hoechst 34580, 405 nm), NPs (561 nm), and zymosan (488 nm) were each represented as a rendered volume. Zymosan was present in phagocytes only. NP and zymosan surfaces were classified as internal or external by their position relative to the membrane surface, and orthogonal xz and yz sections were inspected through every reconstructed cell to confirm that the signal lay beneath the apical membrane rather than on it. These reconstructions establish the intracellular location of the fluorescent signal and were not used for quantification.

### Image analysis and single-cell quantification

Z-stacks from the NP uptake and zymosan phagocytosis experiments were converted to maximum-intensity projections in NIS-Elements and quantified with custom MATLAB scripts (MathWorks). Nuclei were masked from the Hoechst channel and single-cell boundaries from the DiD channel, seeded on the nuclear masks so that each cell mask contained one nucleus; masks outside the expected size range were discarded and segmentation parameters were calibrated to the pixel size of the objective used. Each fluorescent object was assigned to the cell whose mask contained it, and objects outside any cell mask were treated as non-internalized. Intracellular NP fluorescence was measured as the mean 561 nm intensity within each cell mask per unit cell area. Internalized zymosan beads were counted from the 488 nm channel, with clusters converted to bead equivalents by area, and a cell was scored as phagocytic if it contained at least one bead. Because absolute fluorescence varied between imaging sessions, the mean NP intensity of the 0 µg/mL wells was subtracted from all doses within each replicate; statistical tests used the values before subtraction. Cells were classified as NP-positive above the 90th percentile of the 0 µg/mL cells from the same donor and cell type, and NP uptake efficiency is the percentage of NP-positive cells relative to the 0 µg/mL wells. Full segmentation parameters are given in the deposited analysis scripts.

### Statistical analysis

Each biological replicate (N) is an independent experiment using cells from a different donor; technical replicates (n) were averaged to one value per biological replicate before testing. Doses within an experiment came from the same donor, so dose comparisons were paired. N varies between readouts because not every phenotype was obtained from every donor and because imaging fields or wells that failed quality control were excluded. Normality was assessed for every group by the Shapiro-Wilk test. Fluorescence intensities, uptake and phagocytosis efficiencies, viability, bead counts, cell areas, and analyte concentrations were log2-transformed before parametric testing. Hydrodynamic diameter, zeta potential, and secreted MPO were compared on raw values.

Dose comparisons were made by one-way ANOVA with Tukey post-hoc comparisons of all dose pairs, in the repeated-measures form where every donor had a value at every dose and excluding donors who did not; the exception is endothelial area coverage and nuclei counts, where a mixed-effects model fitted by restricted maximum likelihood with donor matching retained donors missing one dose. Where a second factor was present, two-way ANOVA tested both factors, and pairwise comparisons were made within each timepoint because the two endothelial timepoints were run on separate donor sets. Metabolic activity was compared by the Friedman test with Dunn’s multiple comparisons because the values were not normally distributed. Multiplex analytes were compared with the matched 0 µg/mL well from the same donor by paired t-test on log2-transformed concentrations with Benjamini-Hochberg adjustment; analytes at the lower limit of quantification in every well could not be tested. HUVECs and hBMECs were analyzed separately and were not compared with each other.

For NP fluorescence intensity per cell and NP uptake efficiency, the plotted bars are background-subtracted but the statistics were computed before subtraction. To relate zymosan uptake to NP content, cells of each donor and phenotype were pooled across doses and divided into quartiles of single-cell NP intensity, and the mean number of zymosan beads per cell in each quartile was taken as that donor’s value. The test, replicate numbers and corrections for each readout are given in the figure legends, and bar graphs show the mean ± s.e.m. with the mean of individual biological replicates overlaid. A two-sided P value < 0.05 was considered statistically significant. All statistical tests were performed in GraphPad Prism 11.1 (GraphPad Software). Images from the macrophage assays and neutrophil phagocytosis assay and NP uptake assay were processed, quality controlled and reduced to one value per biological replicate in MATLAB R2025b (MathWorks) with custom scripts. All EC image processing was completed in NIS-elements using custom macros. The MATLAB scripts, NIS macros, and the Excel outputs for analysis in Prism are deposited at Zenodo.

## Acknowledgements

Electron microscopy sample preparation and imaging was done at the University of Colorado, Boulder EM Services Core Facility in the MCDB Department, with the technical assistance of facility staff. The authors would like to thank Dr. David Kelaita (Joel L. Kaar and Daniel K. Schwartz labs, University of Colorado Boulder) for training on the Anton Paar Litesizer 500 and assistance in experimental planning. Three-dimensional reconstructions were performed at the University of Colorado Boulder BioFrontiers Institute’s Advanced Light Microscopy Core (RRID: SCR_018302). The Analysis Workstation and the software package Imaris were supported by NIH 1S10RR026680-01A1. We thank the staff of the Core for access and technical support.

## Funding

This work was supported by National Institutes of Health grant R35 GM146737A awarded to LEH, NSF DGE-1746047 to YII, by the Biophysics T32 training grant 5T32GM145437-04 to MR, by the Biofabrication T32 Training Grant 5T32EB034215-03 to LPH, and by the Department of Chemical and Biological Engineering at the University of Colorado Boulder.

## Author contributions

A.I.-R., Y.I.I., J.M.O., and L.E.H. conceived and designed the study. A.I.-R. performed the macrophage experiments, Y.I.I. performed the endothelial cells experiments with assistance from M.R., J.M.O. and J.W.B. performed the neutrophil experiments; E.L.G. performed the neutrophil adhesion experiments. A.I.-R., Y.I.I., and J.M.O. developed the image-analysis and statistical pipelines and analyzed the data. A.I.-R., Y.I.I., J.M.O., and L.E.H. interpreted the results and wrote the manuscript. L.E.H. supervised the project and acquired funding. H.K.W. and L.P.H. contributed to scientific discussion. All authors read and approved the final manuscript.

## Competing interests

The authors declare that they have no competing interests.

## Data and materials availability

Source data are provided with this paper. The MATLAB code and raw data files used to run the statistical tests and to generate the figures are deposited at Zenodo (https://doi.org/10.5281/zenodo.23166506)(^64^).

---

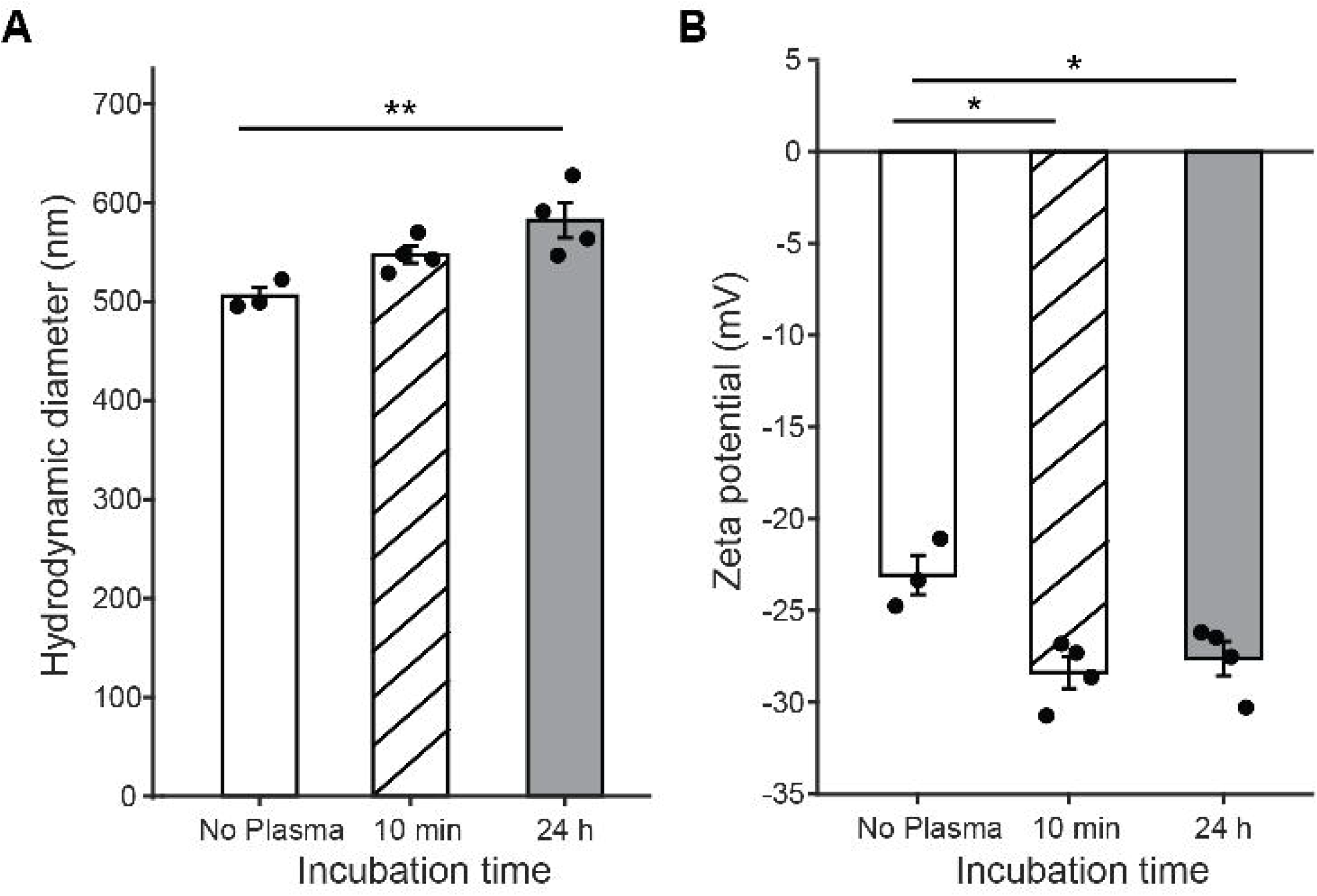

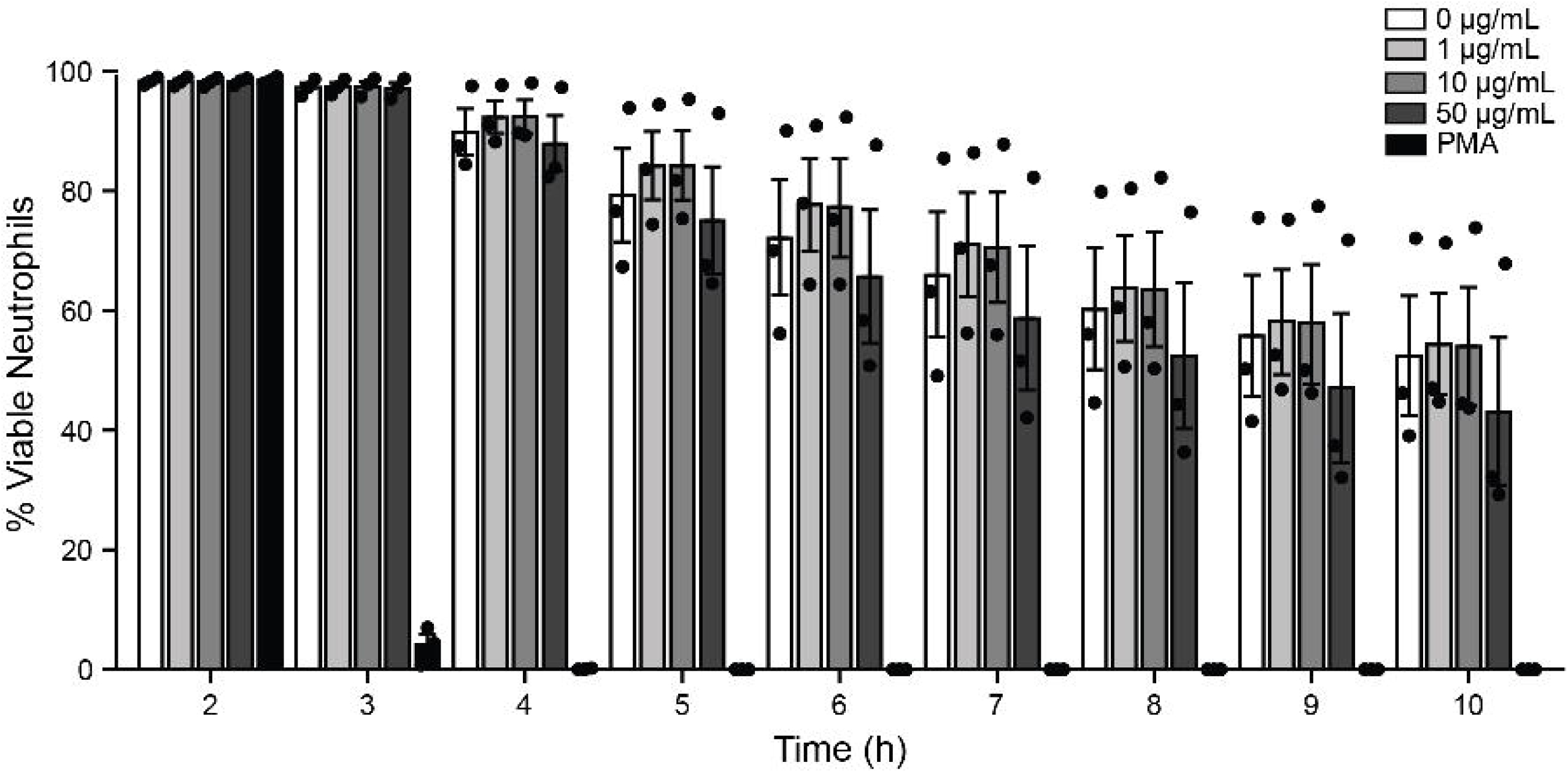

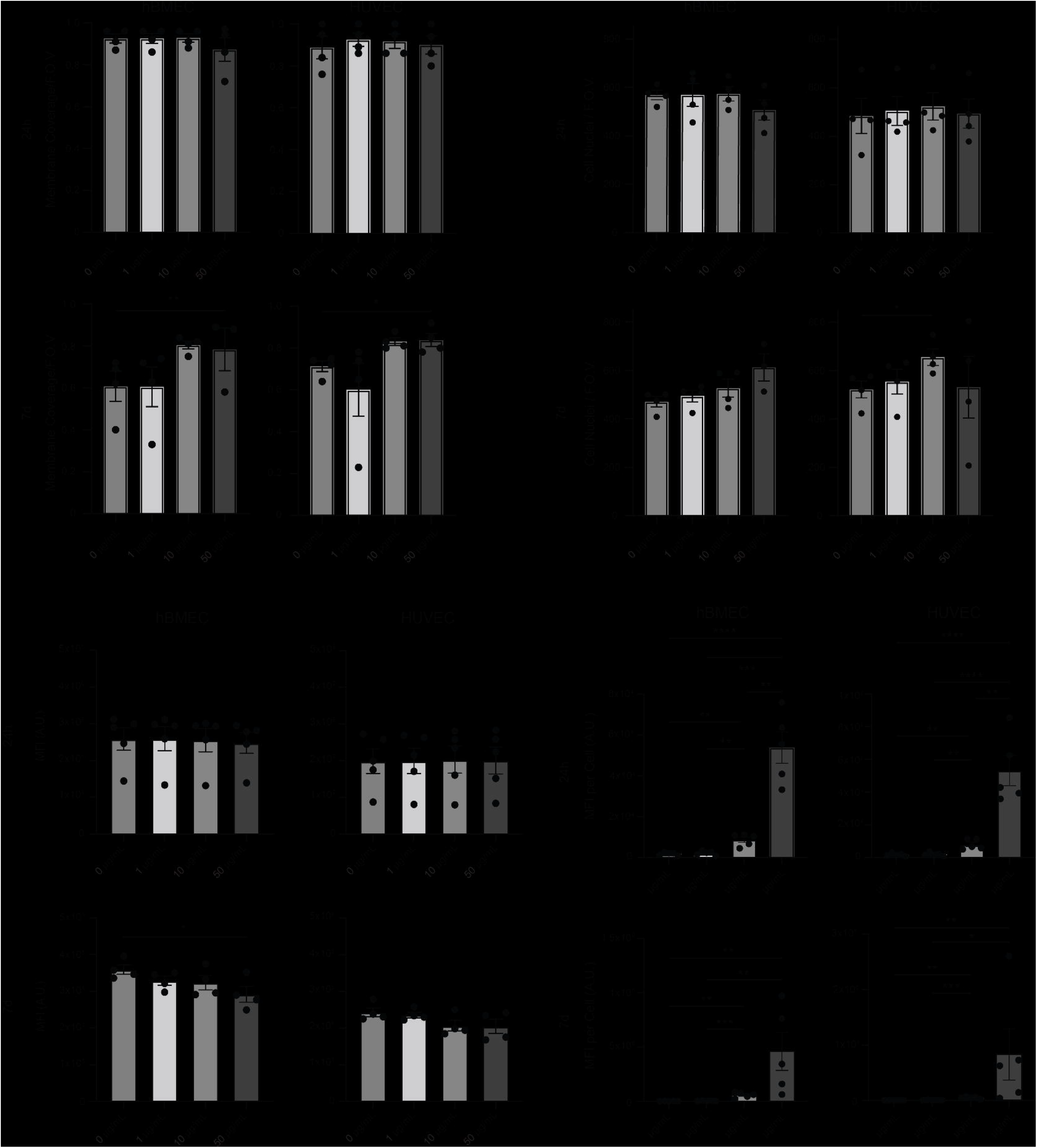

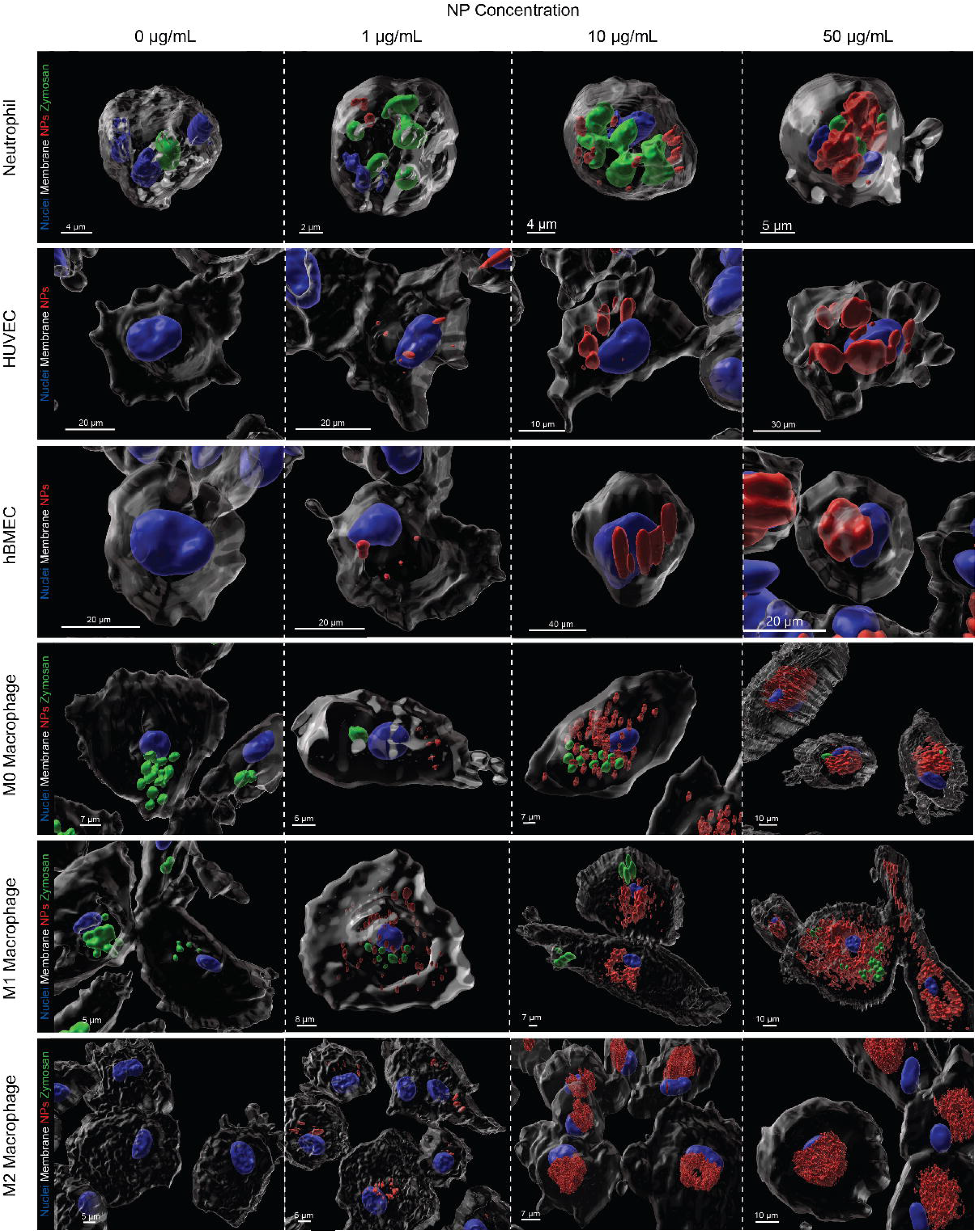

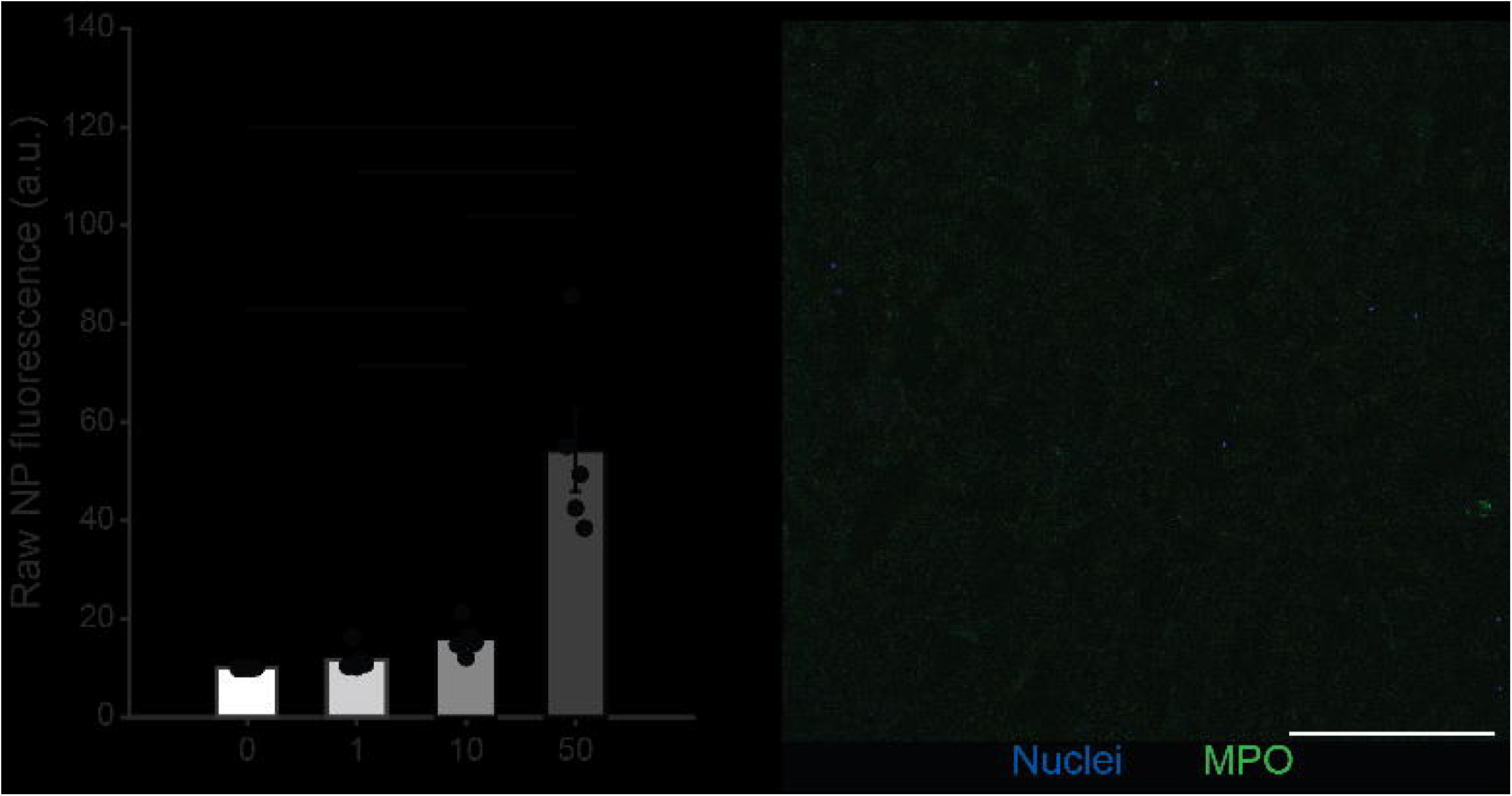

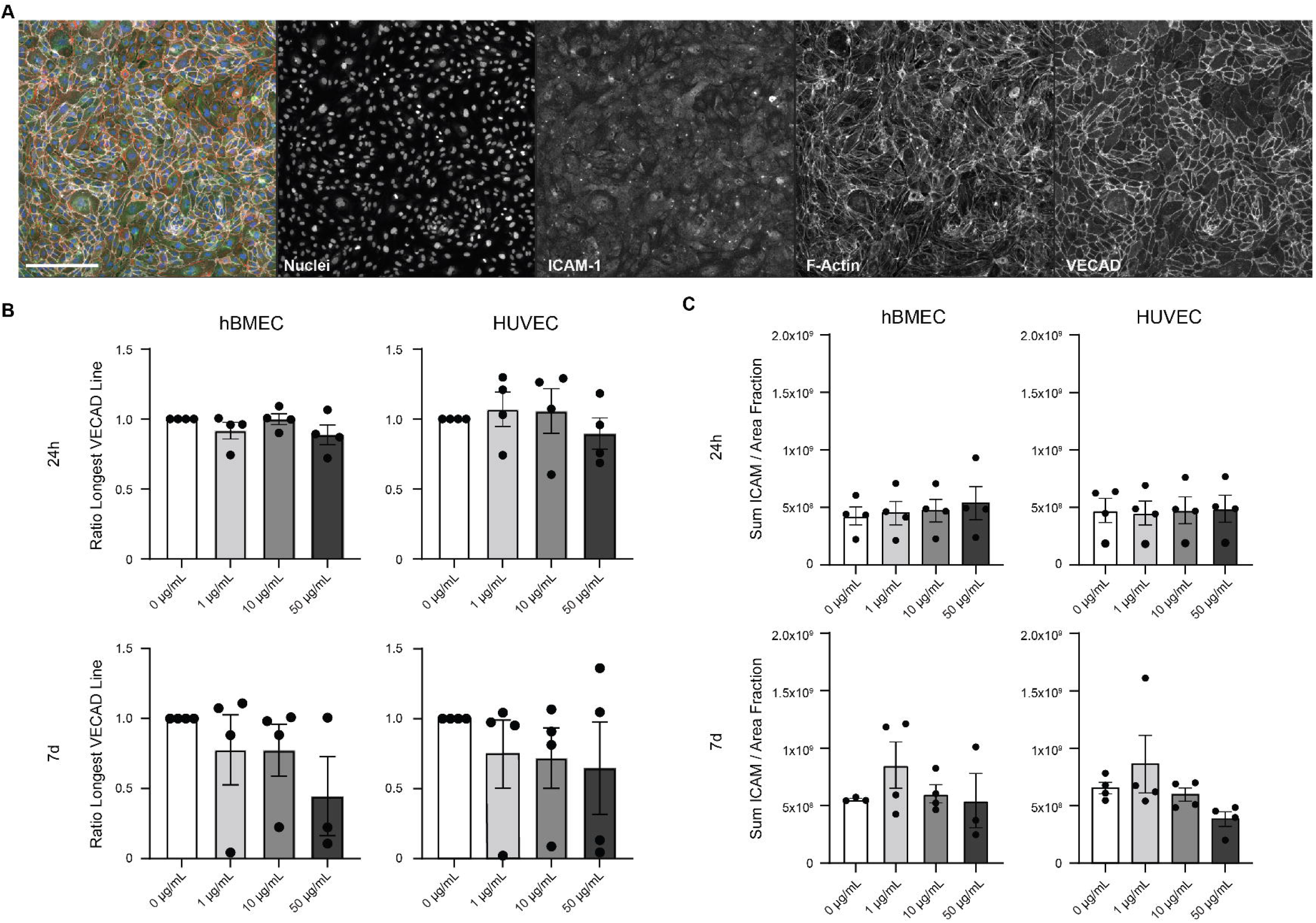

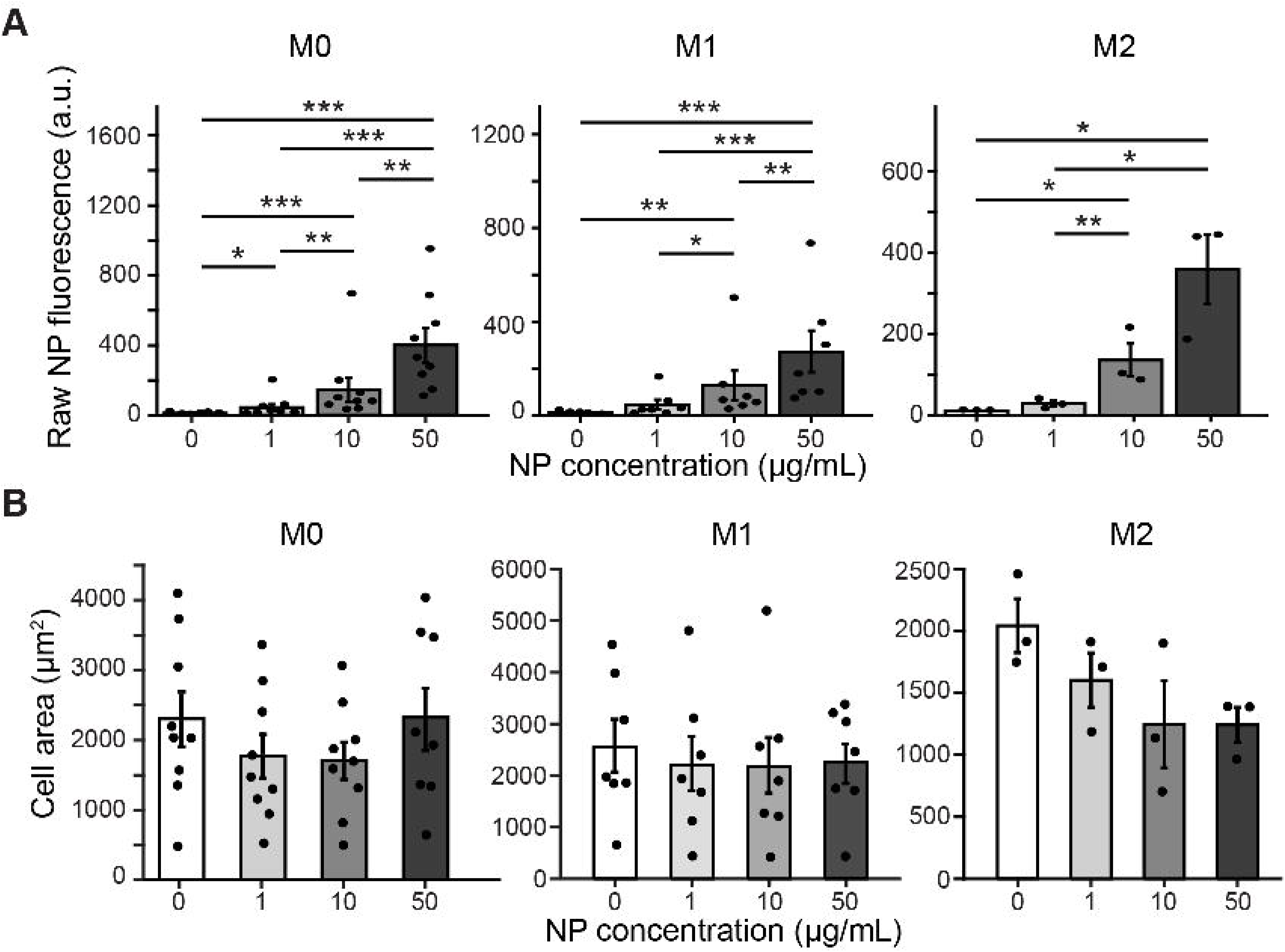

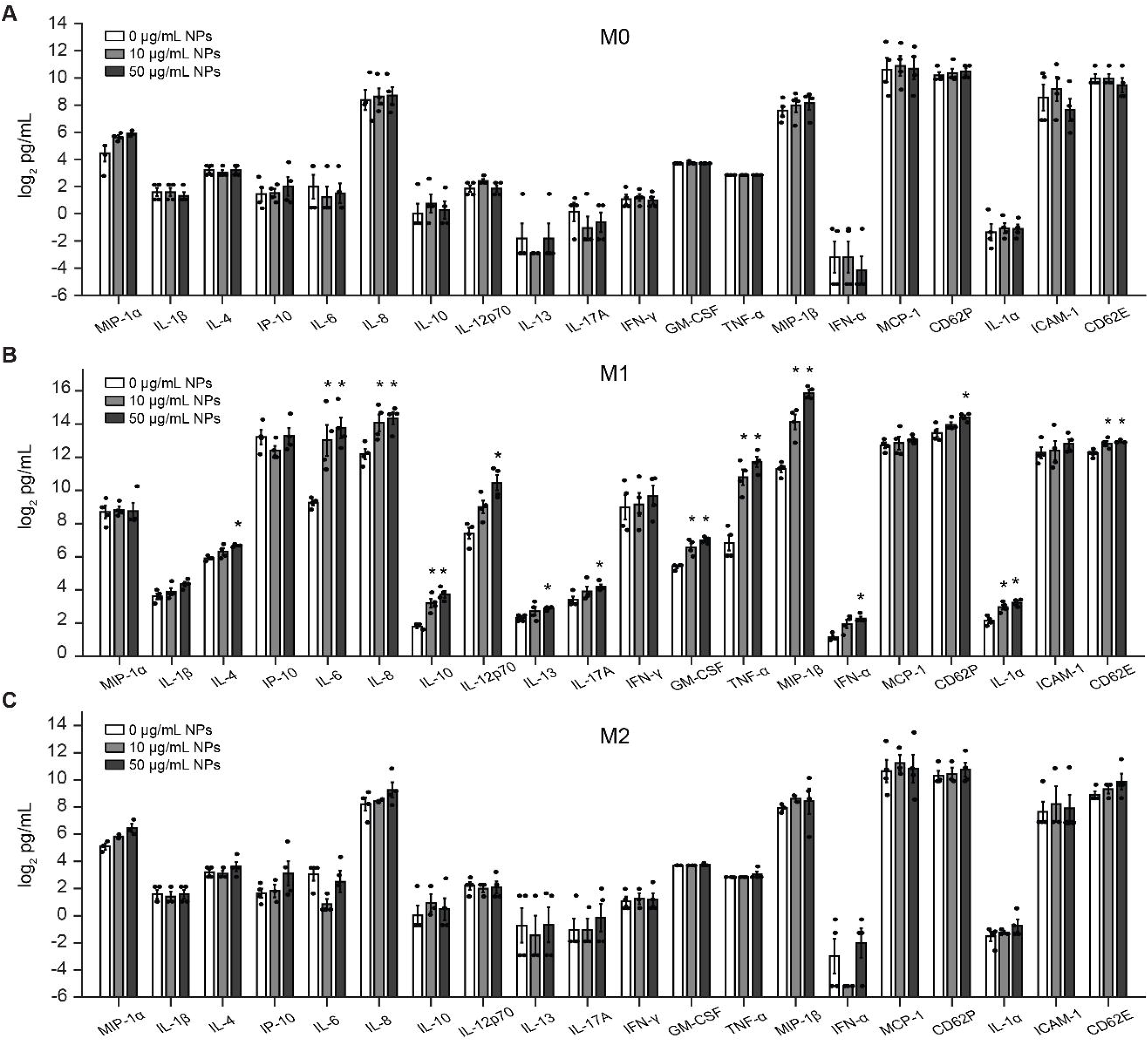

## Notes

### Competing Interest Statement

The authors have declared no competing interest.

