## Supplemental Data for "Cell identity and activation state determine the innate immune response to nanoplastics"

### Supplementary Figure and Figure Legends

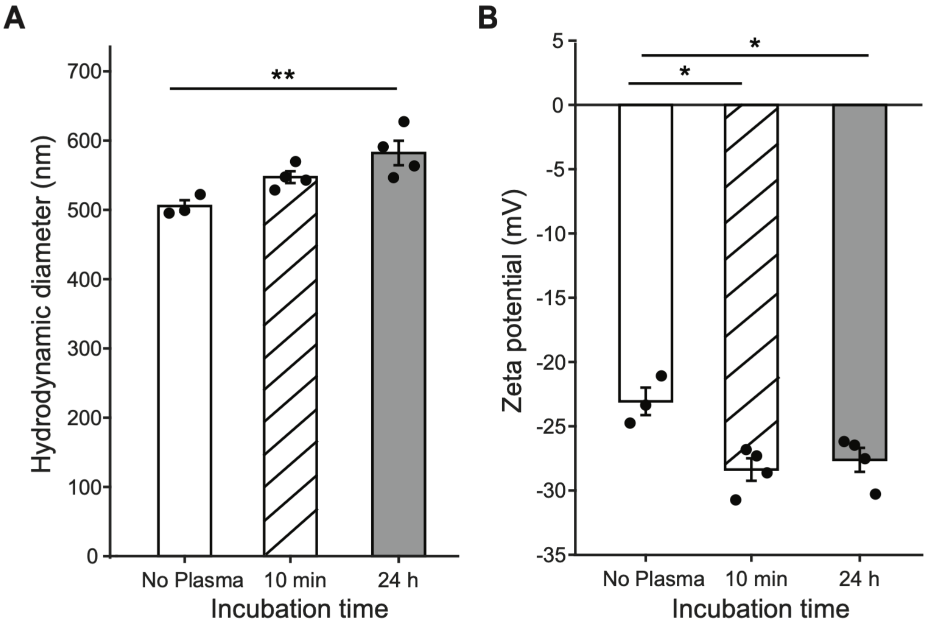


**Supplementary Fig. S1. Fluorescent nanoplastic incubation in human plasma creates a biocorona. A** Fluorescent NP hydrodynamic diameter. **B** Fluorescent NP zeta potential. (**A-B**) One-way ANOVA with Tukey's post-hoc comparisons. N = 4, each biological replicate using plasma from a different donor. Bars show the mean ± s.e.m. with individual biological replicates overlaid. $^*P < 0.05$, $^{**}P < 0.01$. N, biological replicates (donors).

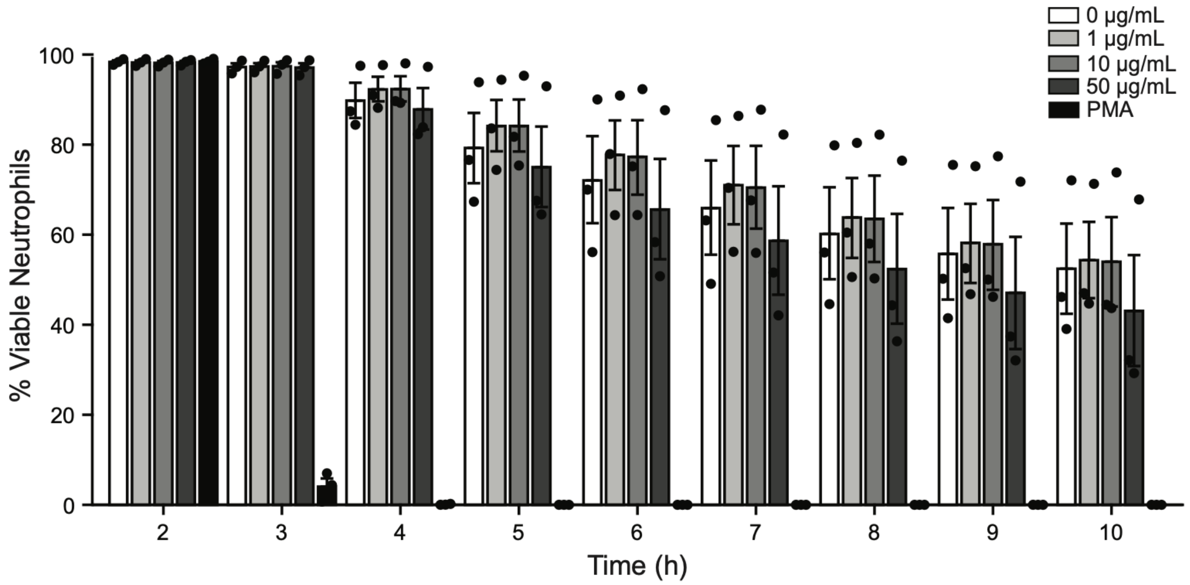


**Supplementary Fig. S2. Neutrophil viability is unaffected by NP concentration up to 10 h.** Neutrophil viability over 10 h after blood draw. Fig. 2B shows the same experiment over the first 6 h. One-way ANOVA with Tukey's post-hoc comparisons. Comparisons between PMA and NP conditions are not shown for clarity. N = 3, n = 6. Points show the mean ± s.e.m. with individual biological replicates overlaid. N, biological replicates (donors); n, technical replicates (fields of view) per biological replicate and condition.

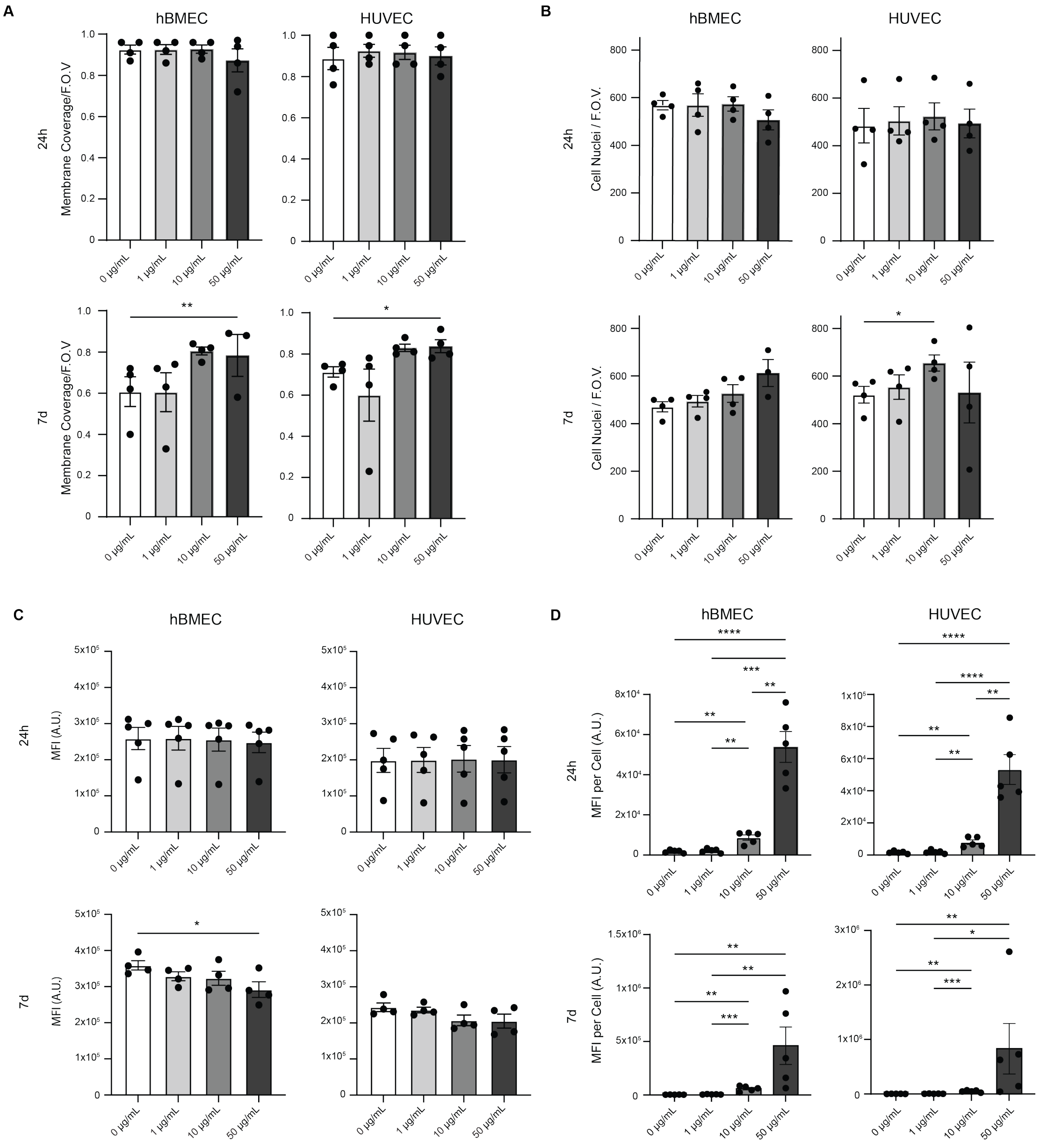


**Supplementary Fig. S3. EC cell count, coverage, and metabolic activity is unchanged even with increased NP accumulation. A** Endothelial cellular coverage reported as fraction of phalloidin coverage per total field of view in hBMECs and HUVECs after 24h or 7d of exposure to 0, 1, 10 or 50 µg/mL non-fluorescent NPs. N=4, n=1. **B** Endothelial cell counts reported as Hoechst+ nuclei per field of view in hBMECs and HUVECs after 24h or 7d of exposure to 0, 1, 10 or 50 µg/mL non-fluorescent NPs. N=4, n=1. (**A-B**) Mixed-effects model (REML) with donor matching and Tukey's post-hoc comparisons. **C** Metabolic activity of ECs measured by alamarBlue fluorescence in hBMECs and HUVECs after 24h or 7d of exposure to 0, 1, 10 or 50 µg/mL non-fluorescent NPs. Friedman test with Dunn's multiple comparisons. N=4-5, n=3. **D**

Summed NP fluorescence intensity divided by number of Hoechst+ nuclei per field of view in hBMECs and HUVECs after 24h or 7d of exposure to 0, 1, 10 or 50 µg/mL fluorescent NPs with no baseline subtraction. Lognormal RM one-way ANOVA with Tukey's post-hoc comparison. N=5, n=9. Bars show the mean ± s.e.m. with individual biological replicates overlaid. *P < 0.05, **P < 0.01, ***P < 0.001, ****P < 0.0001. N, biological replicates (donors); n, technical replicates per biological replicate and condition.

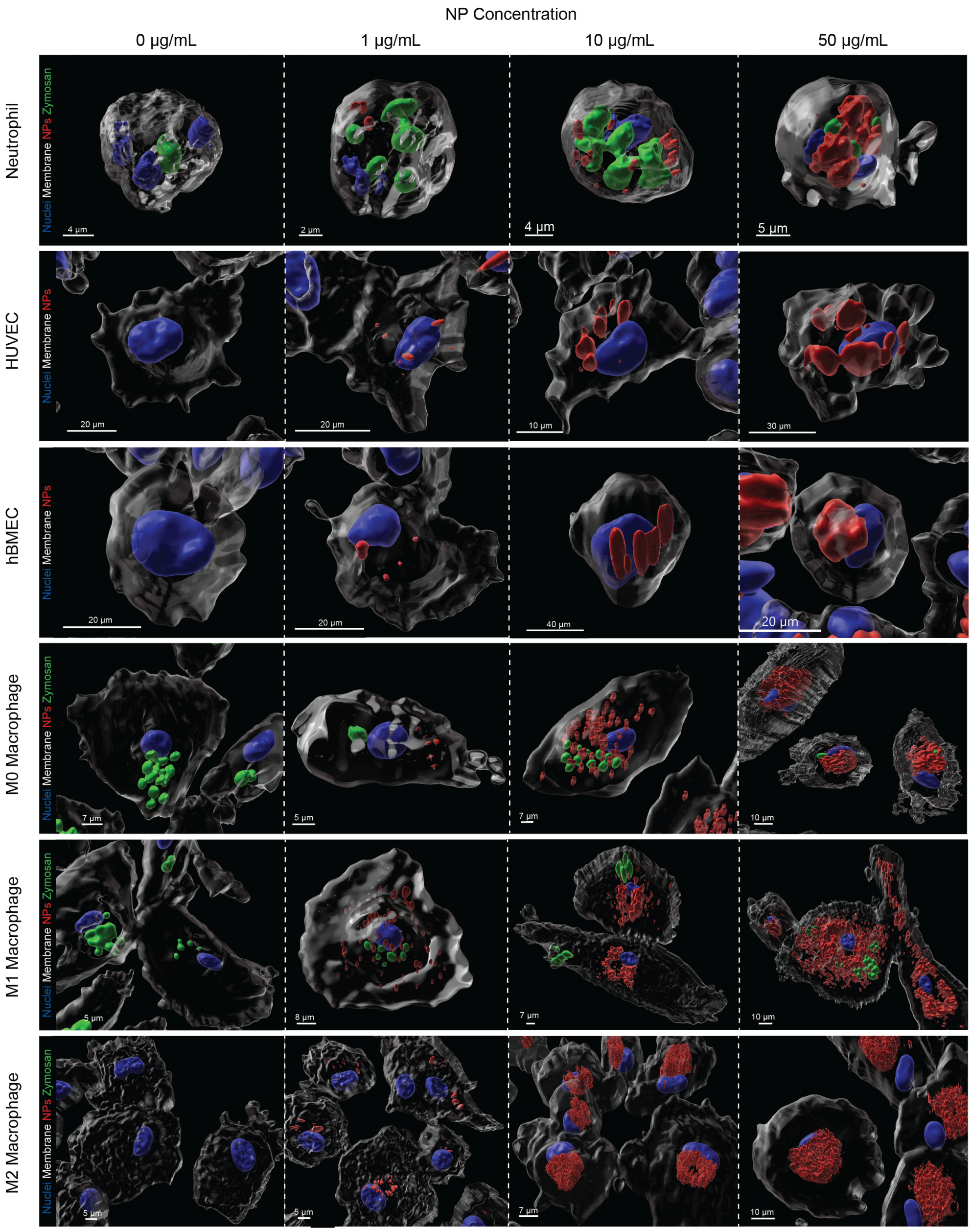


**Supplementary Fig. S4. NPs are internalized by neutrophils, M0, M1 and M2 macrophages, HUVECs and hBMECs.** Three-dimensional Imaris surface renderings of confocal z-stacks of neutrophils, HUVECs and hBMECs, M0, M1 and M2 exposed to 0, 1, 10 or 50 µg/mL biocorona-coated NPs for 24 h. Nuclei (blue), plasma membrane (white) and NPs (red) are shown in all panels; zymosan (green) is shown in the neutrophil and macrophage images. NP and zymosan surfaces were classified as intracellular by their position relative to the rendered plasma membrane. Scale bars, left to right within each row: neutrophils, 4, 2, 4 and 5 µm; HUVECs, 20, 20, 10 and 30 µm;

hBMECs, 20, 20, 40 and 20 μm; M0 macrophages, 7, 5, 7 and 10 μm; M1 macrophages, 5, 8, 7 and 10 μm; M2 macrophages, 5, 5, 7 and 10 μm.

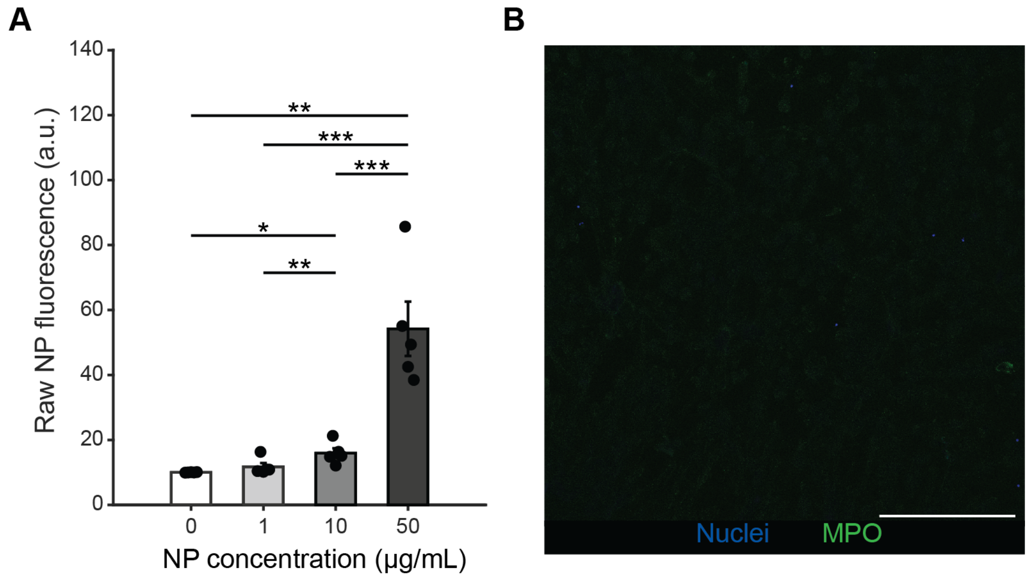


**Supplementary Fig. S5. Neutrophil NP fluorescence without background subtraction and MPO in the PMA control. A** Raw intracellular NP fluorescence without background subtraction; these are the values on which the statistics in Fig. 3D were calculated. One-way repeated-measures ANOVA on log2-transformed values with Tukey's post-hoc comparisons. N = 5, n = 6. **B** Representative image of MPO in the PMA condition, displayed with the same lookup table as the images in Fig. 4D; nuclei (blue) and MPO (green). Scale bar, 250 μm. Bars show the mean ± s.e.m. with individual biological replicates overlaid. *P < 0.05, **P < 0.01, ***P < 0.001. N, biological replicates (donors); n, technical replicates (fields of view) per biological replicate and condition.

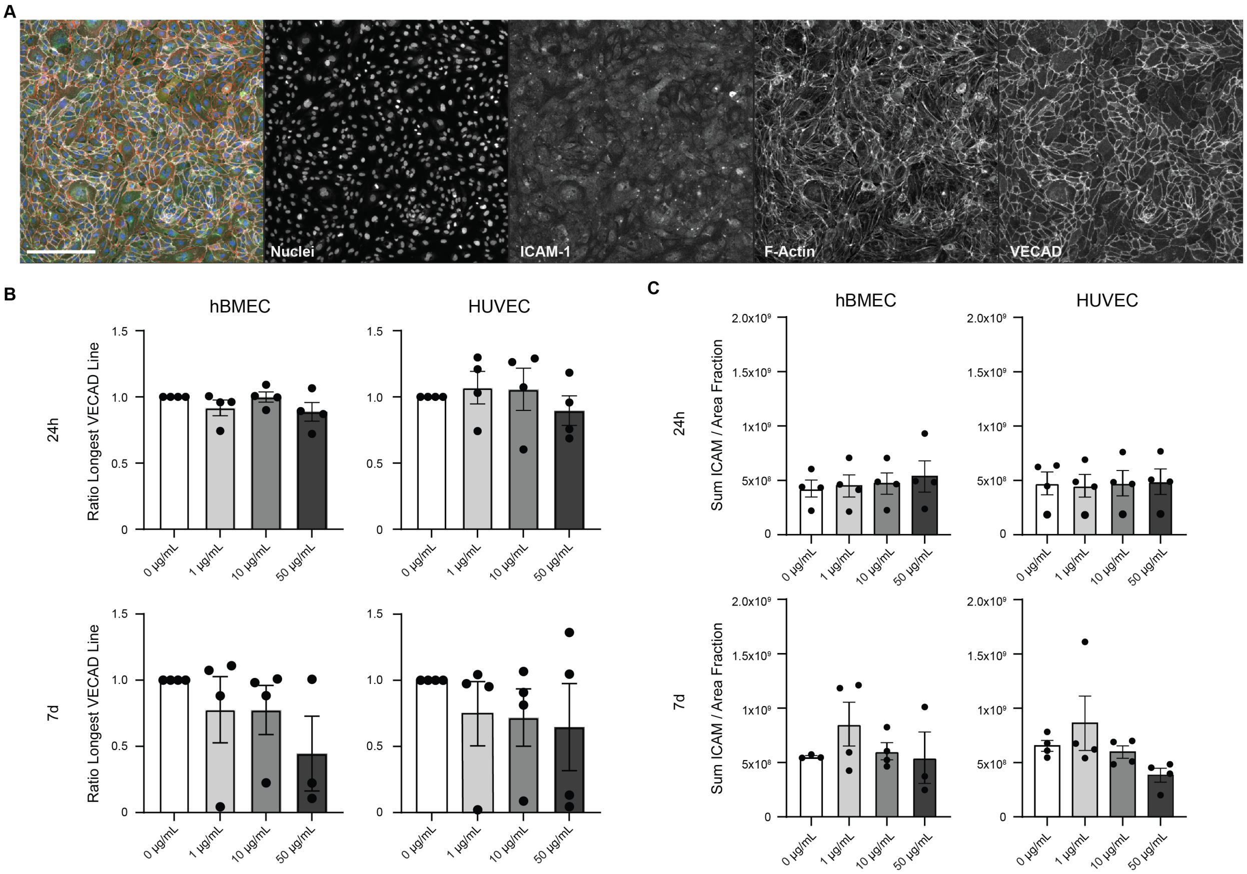


**Supplementary Fig. S6. ECs exhibit no change in VECAD continuity or ICAM-1 expression. A** Representative image of 2D EC staining with Nuclei (blue), ICAM-1 (green), F-Actin (red), and VECAD (white). HUVECs exposed to 10 µg/mL of NPs for 24h shown. Scale bar, 250 µm. **B** VECAD continuity reported as a fraction of the longest continuous VECAD skeleton per NP exposure condition normalized to the 0 µg/mL control. Continuity was quantified in hBMECs and HUVECs after 24h or 7d of exposure to 0, 1, 10 or 50 µg/mL non-fluorescent NPs. Lognormal one-way ANOVA with Tukey's post-hoc comparison. N=4, n=3. **C** Summed intensity of ICAM-1 expressed by ECs normalized by the area coverage determined by phalloidin. ICAM-1 expression was quantified in hBMECs and HUVECs after 24h or 7d of exposure to 0, 1, 10 or 50 µg/mL non-fluorescent NPs. Lognormal one-way ANOVA with Tukey's post-hoc comparison. N=4, n=3. Bars show the mean ± s.e.m. with individual biological replicates overlaid. N, biological replicates (donors); n, technical replicates per biological replicate and condition.

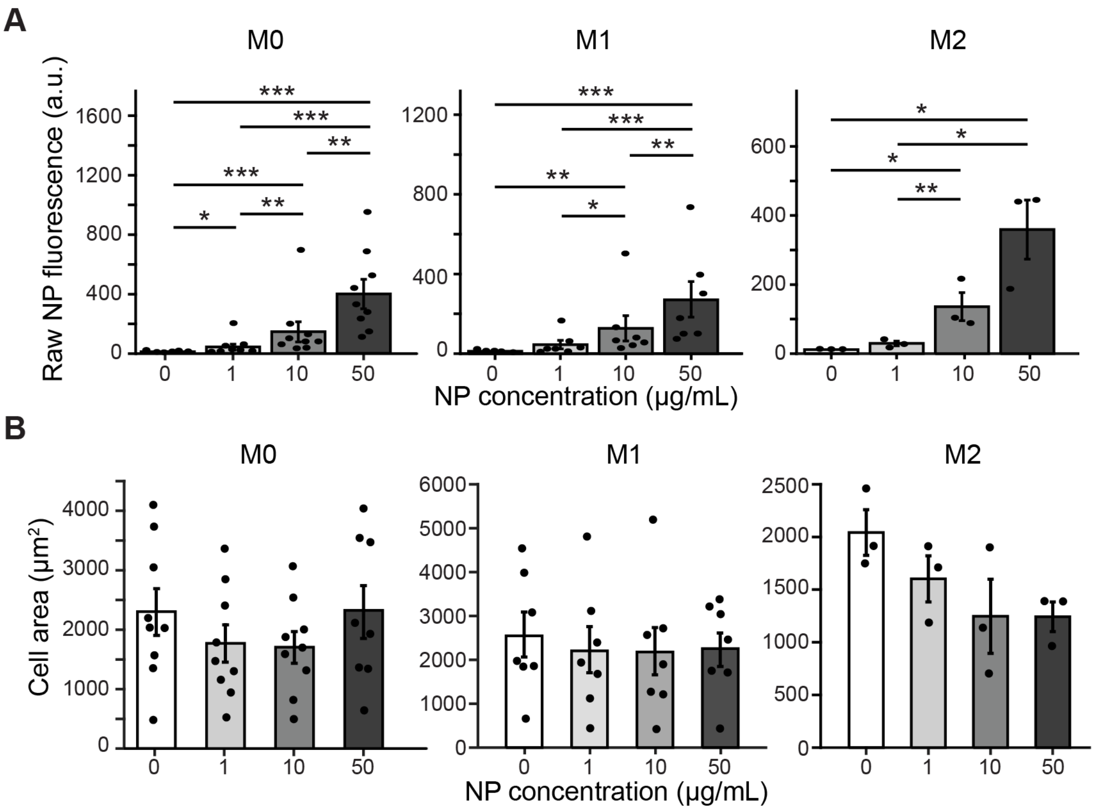


**Supplementary Fig. S7. Macrophage NP fluorescence without background subtraction, cell area and zymosan uptake by NP content. A** NP fluorescence intensity per cell without background subtraction in M0, M1 and M2 macrophages; these are the values on which the statistics in Fig. 3G were calculated. **B** Mean macrophage area. (**A** and **B**) One-way repeated-measures ANOVA on log2-transformed values with Tukey's post-hoc comparisons; no comparison reached significance in B. N = 9 (M0; 8 at 50 μg/mL), 7 (M1) and 3 (M2), n = 1–5. Bars show the mean ± s.e.m. with individual biological replicates overlaid. $^{*}P < 0.05$, $^{**}P < 0.01$, $^{***}P < 0.001$. N, biological replicates (donors); n, technical replicates per biological replicate and condition.

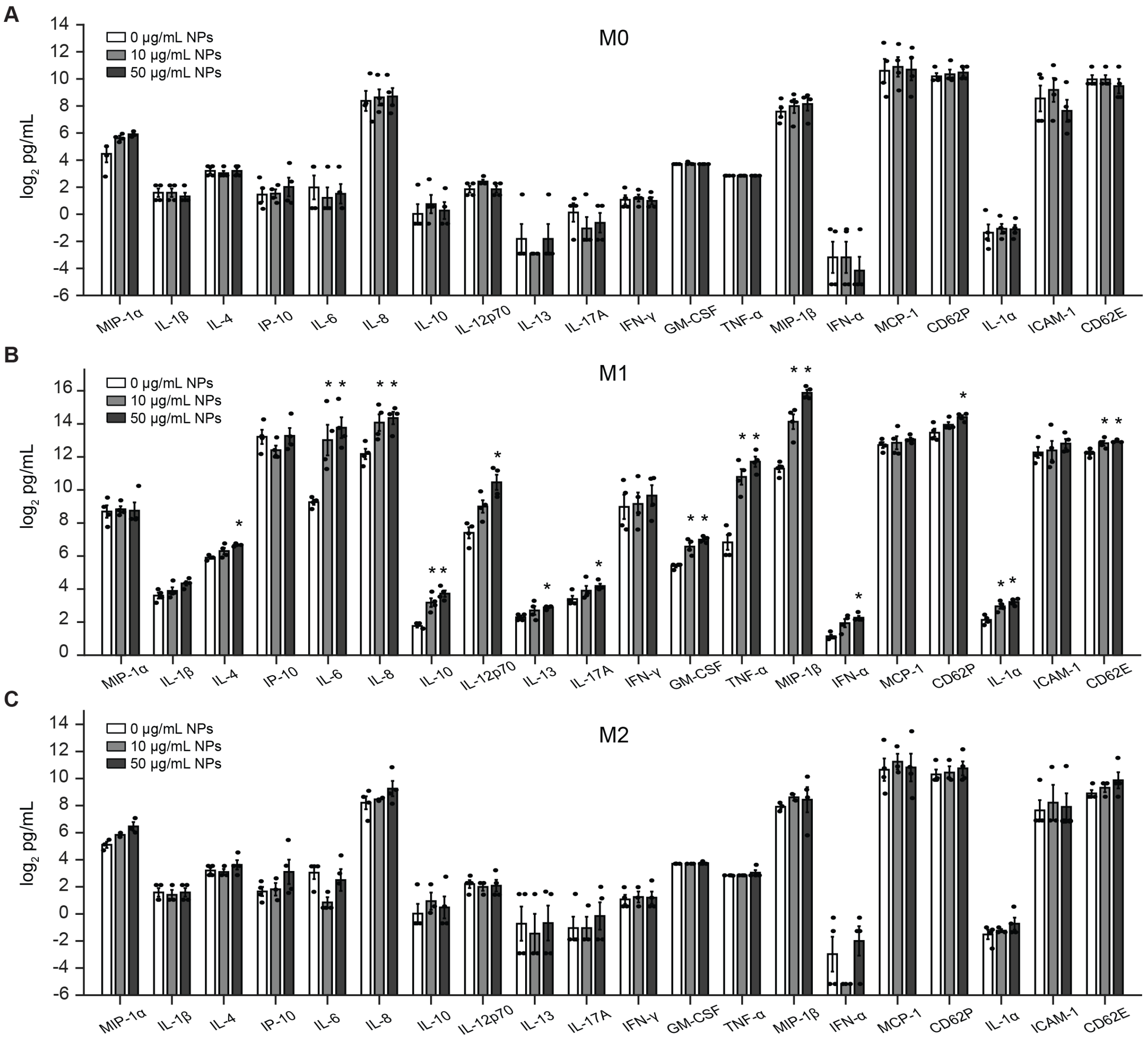


**Supplementary Fig. S8. NP exposure changes cytokine secretion in M1 macrophages but not in M0 or M2 macrophages. A** to **C** Concentrations of all 20 cytokines, chemokines and soluble adhesion molecules in conditioned media of M0 (**A**), M1 (**B**) and M2 (**C**) macrophages after 24 h exposure to 0, 10 or 50 μg/mL non-fluorescent NPs, quantified by multiplex immunoassay (MAGPIX, 20-plex) and shown as log2 pg/mL. Panel B shows the same data as Fig. 5E. Values below the lowest standard were assigned the lower limit of quantification for that analyte. Each concentration was compared with the matched 0 μg/mL well from the same donor by two-sided paired t-test on log2-transformed concentrations, with Benjamini-Hochberg correction across all 20 analytes and both concentrations, separately for each phenotype; no comparison reached significance in M0 or M2 macrophages. N = 3–4 (M0), 4 (M1) and 2–4 (M2), n = 1; wells and channels excluded before analysis are listed in the deposited data. Bars show the mean ± s.e.m. of log2 concentrations with individual biological replicates overlaid. *Adjusted $P < 0.05$. N, biological replicates (donors); n, technical replicates per biological replicate and condition.
